## Supplementary material for "Reconciliation and evolution of *Penicillium rubens* Genome-Scale Metabolic Networks – What about specialised metabolism?"

### *Table of Contents: Supplementary Tables*

**Table S1: Functional annotation details.**

| *Penicillium rubens* Wisconsin 54-1255[1] | |
|---|---|
| **Nature of the original data** | |
| Number of sequences | 12,556 |
| Average sequence length (residue counts) | 1,343 |
| Minimum sequence length (nucleotides) | 48 |
| Maximum sequence length (nucleotides) | 21,864 |
| Average sequence length (residue counts) | 447 |
| Minimum sequence length (amino acids) | 15 |
| Maximum sequence length (amino acids) | 7,287 |
| **Protein domain research (hmmscan[2])** | |
| Total number of protein domains detected | 19,589 |
| Number of different protein domains detected | 4,439 |
| Number of sequences annotated by at least one protein domain | 8,607 (69%) |
| Number of sequences involved in the sub-network annotation | 3,296 |
| Number of sequences involved in the draft | 5,193 |
| **Nature of proteins** | |
| Sequences with transmembrane helices (TMHMM[3]) | 2,283 (18.1818%) |
| Sequences with signal peptide (SignalP[4]) | 921 (7.3%) |
| **Gene Ontology Term (GOT[5])** | |
| Number of sequences annotated by at least one GOT (Trinotate[6]) | 8,042 (64%) |
| Total number of GOTs detected | 429,742 |
| Number of different GOTs detected | 12,712 |
| Number of sequences involved in the sub-network annotation | 3,370 |
| Number of sequences involved in the draft | 5,125 |
| **Kyoto Encyclopaedia of Genes and Genomes (KEGG[7])** | |
| Number of sequences annotated by at least one KO (Trinotate[6]) | 4,247 (34%) |
| Number of sequences annotated by at least one KO (KAAS[8]) | 4,082 (33%) |
| Number of sequences annotated by at least one KO (KEGG[7] – genes pcs) | 3,626 (29%) |
| Number of sequences annotated by at least one KO (consensus) | 4,991 (40%) |
| The total number of KEGG detected | 5,105 |
| Number of different KEGG detected | 3,664 |
| **Enzyme Commission numbers (EC)** | |
| Number of sequences annotated at least by one EC (Trinotate[6]) | 1,937 (15%) |
| Number of sequences annotated at least by one EC (KAAS[8]) | 1,849 (15%) |
| Number of sequences annotated at least by one EC (KEGG[7] – pcs genes) | 1,625 (13%) |
| Number of sequences annotated by at least one EC (consensus) | 2,335 (19%) |
| Total number of EC detected | 2,603 |
| Number of different EC detected | 1,206 |
| Number of sequences involved in the sub-network annotation | 2,321 |
| Number of sequences involved in the draft | 2,333 |

### *References*

**Table S2: Features and results of the seven templates used for the reconstruction of the intermediate sub-network from the orthology search.**

| **Organism** | ***A. thaliana*** | ***A. nidulans*** | ***A. niger*** | ***N. crassa*** | ***P. rubens* species** | ***S. japonica*** | ***S. pombe*** |
|---|---|---|---|---|---|---|---|
| **Features of the networks used** | | | | | | | |
| Publication | de Oliveira Dal'Molin *et al.*[1]. | Pitkänen *et al.*[2] Castillo *et al.*[3] | Pitkänen *et al.*[2] Castillo *et al.*[3] | Dreyfuss *et al.*[4] | Pitkänen *et al.*[2] Castillo *et al.*[3] | Nègre *et al.*[5] | Pitkänen *et al.*[2] Castillo *et al.*[3] |
| Main database | KEGG[6] | KEGG[6] | KEGG[6] | Undefined | KEGG[6] | MetaCyc[7] | KEGG[6] |
| Date | 2010 | 2016 | 2016 | 2014 | 2016 | 2018 | 2016 |
| Number of reactions | 1,510 | 3,689 | 3,703 | 1,392 | 3,666 | 3,286 | 3,158 |
| Number of Metabolites | 1,514 | 3,196 | 3,206 | 736 | 3,176 | 3,231 | 2,749 |
| Number of genes \| proteins | 2,330 | 1,279 | 1,299 | 836 | 1,352 | 5,017 | 810 |
| **OrthoFinder[8] Results and associated reaction** | | | | | | | |
| Number of *P. rubens* orthologous genes | 866 | 1,818 | 1,620 | 1,104 | 2,368 | 2,329 | 1,114 |
| Number of reactions detected | 949 | 2,912 | 2,680 | 1,008 | 2,886 | 2,487 | 2,508 |
| **Integration to the draft** | | | | | | | |
| Number of reactions that map to MetaCyc | 441 | 1,434 | 1,307 | 462 | 1,425 | 2,167 | 1,222 |
| Number of genes integrated into the draft | 586 | 1,347 | 1,182 | 667 | 1,709 | 2,188 | 858 |

### *References*

**Table S3: List of reactions involved in the synthesis of roquefortines and meleagrin added to the reconstruction (MetaCyc identifiers).**

| Pathway (MetaCyc Identifiers[1]) | Reaction (MetaCyc Identifiers[1]) | Gene |
|---|---|---|
| **PWY-7609** | RXN-16124 | roqA (rds) - *Pc21g15480* |
| **PWY-7609** | RXN-16125 | roqR - *Pc21g15470* |
| **PWY-7609** | RXN-16126 | roqD (rpt) - *Pc21g15430* |
| **PWY-7609** | RXN-16127 | roqD (rpt) - *Pc21g15430* |
| **PWY-7609** | RXN-16128 | roqR - *Pc21g15470* |
| **PWY-7609** | RXN-16140 | roqM - *Pc21g15460* |
| **PWY-7609** | RXN-16141 | roqM - *Pc21g15460* |
| **PWY-7609** | RXN-16142 | roqM - *Pc21g15460* |
| **PWY-7609** | RXN-16143 | roqO - *Pc21g15450* |
| **PWY-7609** | RXN-16144 | roqO - *Pc21g15450* |
| **PWY-7609** | RXN-16145 | roqN (gmt) - *Pc21g15440* |
| **PWY-7609** | RXN-16146 | roqN (gmt) - *Pc21g15440* |
| **PWY-7609** | RXN-16147 | roqO - *Pc21g15450* |

### *Reference*

(1) Caspi, R.; Billington, R.; Keseler, I. M.; Kothari, A.; Krummenacker, M.; Midford, P. E.; Ong, W. K.; Paley, S.; Subhraveti, P.; Karp, P. D. The MetaCyc Database of Metabolic Pathways and Enzymes - a 2019 Update. Nucleic Acids Res. **2020**, 48 (D1), D445–D453. https://doi.org/10.1093/nar/gkz862.

**Table S4: Lists of reactions involved in penicillins biosynthesis. GPRs associations in agreement with the literature are shown in green.**

| MetaCyc Identifier[1] | | Presence in the reconstruction | Expected gene | Genes in the reconstruction |
|---|---|---|---|---|
| **Pathways** | **Reactions** | | | |
| **PWY-5629** | 6.3.2.26-RXN | Orthology *S. pombe* | ACVS - *Pc21g21390* | *Pc22g06310* |
| | | Orthology *S. pombe* | | *Pc22g06680* |
| | | Orthologie *A. nidulans* | | *Pc21g01710* |
| | | Orthology PENCH | | *Pc21g09220* |
| | | Orthology PENCH | | *Pc16g07090* |
| | | Orthology PENCH | | *Pc16g06630* |
| | | Orthology PENCH | | *Pc21g20360* |
| | | Orthology PENCH | | *Pc21g05310* |
| | | Orthology PENCH | | *Pc12g14890* |
| | | Orthology PENCH | | *Pc22g21580* |
| **PWY-5629** | 1.21.3.1-RXN | Orthology *S. pombe* | IPNS - *Pc21g21380* | *Pc13g09880* |
| | | Orthology *A. nidulans* & Annotation (Ec-number) | | ***Pc21g21380*** |
| | | Orthology PENCH | | *Pc21g08180* |
| | | Orthology *A. niger* | | *Pc21g06000* |
| | | Orthology *A. niger* | | *Pc22g05480* |
| | | Orthology *S. pombe* | | *Pc15g01820* |
| **PWY-7716** | RXN-10819 | Manual (from Prubens) | phl - *Pc22g14900* | ***Pc22g14900*** |
| **PWY-7716** | RXN-17061 | Annotation (Ec-number) | PenDE - *Pc21g21370* | ***Pc21g21370*** |
| | | Annotation (GOT) | | *Pc13g09140* |
| **PWY-7716** | RXN-17062 | Annotation (Ec-number) | PenDE - *Pc21g21370* | ***Pc21g21370*** |
| | | Annotation (GOT) | | *Pc13g09140* |
| **PWY-7716** | RXN-17100 | - | PenDE - *Pc21g21370* | - |
| **PWY-5630** | RXN-8809 | Annotation (Ec-number) | PenDE - *Pc21g21370* | ***Pc21g21370*** |
| | | Annotation (GOT) | | *Pc13g09140* |

| ***Reference*** |
| --- |

(1) Caspi, R.; Billington, R.; Keseler, I. M.; Kothari, A.; Krummenacker, M.; Midford, P. E.; Ong, W. K.; Paley, S.; Subhraveti, P.; Karp, P. D. The MetaCyc Database of Metabolic Pathways and Enzymes - a 2019 Update. Nucleic Acids Res. **2020**, 48 (D1), D445–D453. https://doi.org/10.1093/nar/gkz862.

**Table S5: Identification comparison of BGCs between antiSMASH and fungiSMASH.**

| Type | Name | AntiSMASH | FungiSMASH | MetaCyc ID |
|---|---|---|---|---|
| Siderophores | Pistillarin | 0 | 0 | - |
| Siderophores | Coprogen | 0 | 0 | CPD0-2262 |
| Siderophores | Fusarinine | 0 | 0 | CPD-19128 |
| Siderophores | Ferrichrome | 0 | 0 | CPD0-2241 |
| NRPS | Aspercryptins | 1 | 0 | - |
| NRPS | Nidulanin A | 1 | 1 | - |
| NRPS | Benzylpenicillin (penicillin G) | 1 | 1 | PENICILLIN-G |
| NRPS | Isopenicillin N | 1 | 0 | ISOPENICILLIN-N |
| NRPS | Phenoxymethylpenicillin (penicillin V) | 1 | 0 | CPD-9196 |
| NRPS | δ-(L-α-aminoadipyl)-L-cysteine-D-valine (ACV) | 1 | 0 | N-5S-5-AMINO-5-CARBOXYPENTANOYL-L-CY |
| Terpene | PR-toxin | 1 | 0 | CPD-13303 |
| Terpene | Squalestatin S1 | 1 | 1 | - |
| T1PKS | Andrastin | 0 | 0 | - |
| T1PKS | Naphthopyrone | 0 | 1 | - |
| T1PKS | Chrysoxanthone A | 0 | 1 | - |
| T1PKS | Chrysoxanthone B | 0 | 1 | - |
| T1PKS | Chrysoxanthone C | 0 | 1 | - |
| T1PKS | Depudecin | 0 | 1 | - |
| T1PKS | Neurosporin A | 1 | 0 | - |
| T1PKS | ACT-Toxin II | 1 | 0 | - |
| T1PKS | Patulin | 1 | 1 | CPD-16726 |
| T1PKS | Melanin | 1 | 0 | MELANIN |
| T1PKS | Sorbicillin | 1 | 1 | - |
| T1PKS | Yanuthone D | 1 | 1 | - |
| T1PKS,NRPS | Chrysogine | 1 | 1 | - |
| T1PKS,NRPS | Dimethylcoprogen | 1 | 1 | - |
| T1PKS,NRPS | Chaetoglobosins | 1 | 0 | - |
| T1PKS,NRPS | NG-391 | 1 | 0 | - |
| T1PKS,NRPS | Dehydrohistidyltryptophanyldiketopiperazine (DHTD) | 1 | 1 | CPD-17379 |
| T1PKS,NRPS | Glandicoline A | 1 | 1 | CPD-17389 |
| T1PKS,NRPS | Glandicoline B | 1 | 1 | CPD-17390 |
| T1PKS,NRPS | Histidyltryptophanyldiketopiperazine (HTD) | 1 | 1 | CPD-17378 |
| T1PKS,NRPS | Meleagrine | 1 | 1 | CPD-17391 |
| T1PKS,NRPS | Roquefortine C | 1 | 1 | CPD-17381 |
| T1PKS,NRPS | Roquefortine D | 1 | 1 | CPD-17380 |

| Biosynthetic Genes Clusters (BGCs) | antiSMASH[1] | fungiSMASH[1] |
|---|---|---|
| **Numbers of BGCs (identified compounds)** | 45 (16) | 42 (12) |
| **Terpenes (identified compounds)** | 11 (2) | 5 (1) |
| **NRPS (identified compounds)** | 21 (3) | 19 (2) |
| **T1PKS (identified compounds)** | 7 (6) | 14 (6) |
| **T1PKS, NRPS (identified compounds)** | 5 (5) | 4 (3) |
| **Siderophore (identified compounds)** | 1 (0) | - |

List of natural products known to occur in *Penicillium rubens* and identified by the SMASH[1] tool suite. Slight differences exist between antiSMASH and fungiSMASH results (noted 1 for presence and 0 for absence). The colours represent the metabolites according to their type (Siderophore, NRPS, Terpene, T1PKS and T1PKS, NRPS) and then their belonging to the same Biosynthetic Gene Cluster (BGC). More than half of these compounds are “orphans” since they are not listed in the MetaCyc[2] database. The identifiers highlighted in red are currently unusable because they do not have sufficiently precise information for their production or consumption (*e.g.* lack of reactions or too generic reactions). The BGC for patulin, written in red, is incomplete in *P. rubens*.

### *References*

(1) Blin, K.; Shaw, S.; Steinke, K.; Villebro, R.; Ziemert, N.; Lee, S. Y.; Medema, M. H.; Weber, T. antiSMASH 5.0: Updates to the Secondary Metabolite Genome Mining Pipeline. Nucleic Acids Res. **2019**, 47 (W1), W81–W87. https://doi.org/10.1093/nar/gkz310.

(2) Caspi, R.; Billington, R.; Keseler, I. M.; Kothari, A.; Krummenacker, M.; Midford, P. E.; Ong, W. K.; Paley, S.; Subhraveti, P.; Karp, P. D. The MetaCyc Database of Metabolic Pathways and Enzymes - a 2019 Update. Nucleic Acids Res. **2020**, 48 (D1), D445–D453. https://doi.org/10.1093/nar/gkz862.

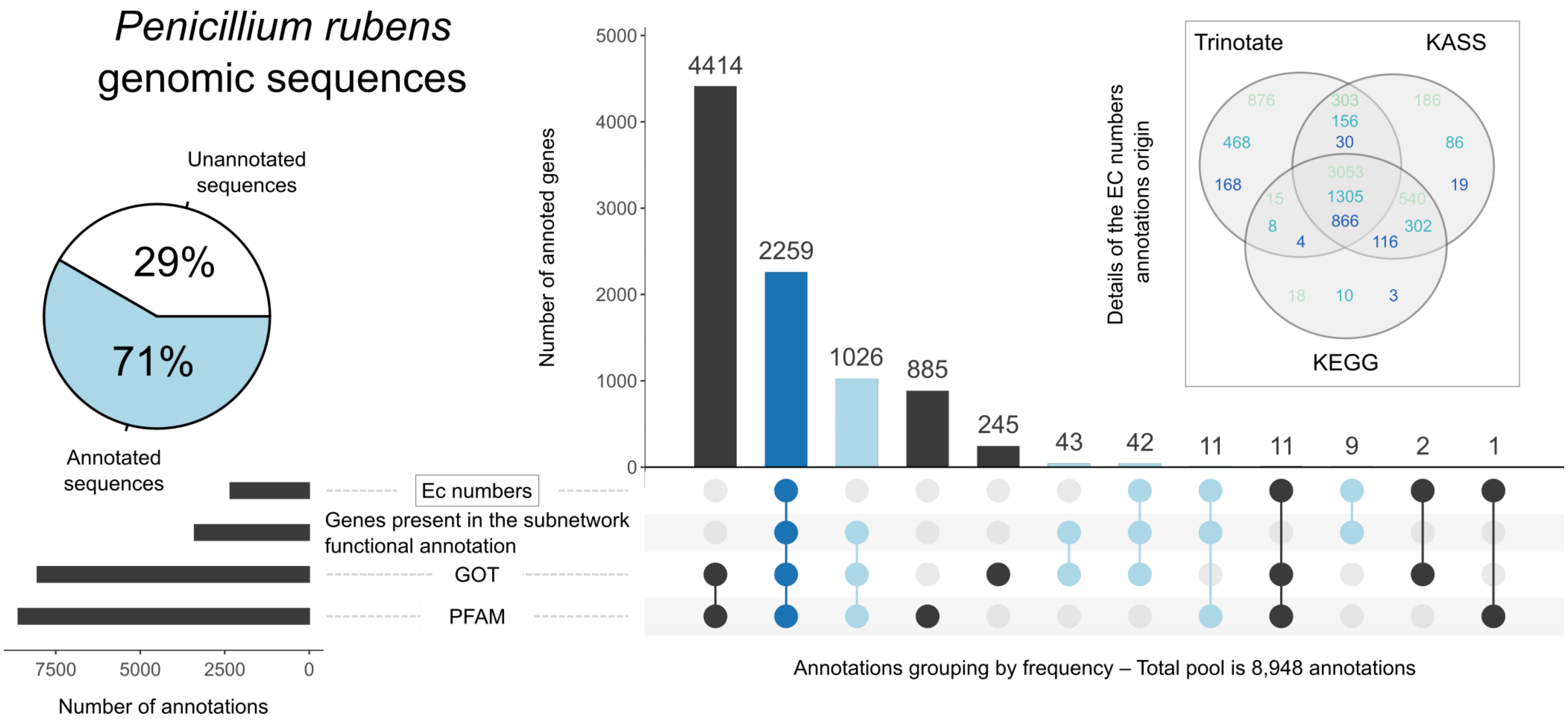

**Figure S1: Functional Annotation Overview of *Penicillium rubens*.** Upset plot illustrating the distribution of three types of annotation data: EC numbers, gene ontology terms (GOT),[1] and protein domains (PFAM)[2] for the 8,948 annotated sequences. Among these genes, 3,390 support at least one reaction within the intermediate functional annotation subnetwork. These genes are highlighted in blue. Specifically, dark blue (■) represents the 2,259 sequences present in this network and annotated with at least one EC number, one GOT, and one PFAM domain, while light blue (■) represents the set of genes annotated with one or two sources. It is worth noting that the vast majority of genes associated with an EC number are found in the draft. The set of 14 genes not included thus constitutes a group of candidates to be explored preferentially for the future completion of the network. Venn diagram representing the impact of the different approaches in obtaining genes annotated with a KO identifier[3] (■), those subsequently annotated with an EC number (■), and the nature of the detected EC numbers (■). Of the 4,501 genes annotated with a KO identifier, 2,343 are associated with an EC number (ratio of about 1:2). 68% of the KOs and 56% of the EC numbers are common to all three approaches. 19% of the KOs and 20% of the EC numbers are provided exclusively by the annotation performed *via* Trinotate[4]. Notably, Trinotate and KAAS effectively retrieve almost all the information contained in the KEGG database.[5] Figure generated with the R package UpSetR (v1.4.0).[6]

### *References*

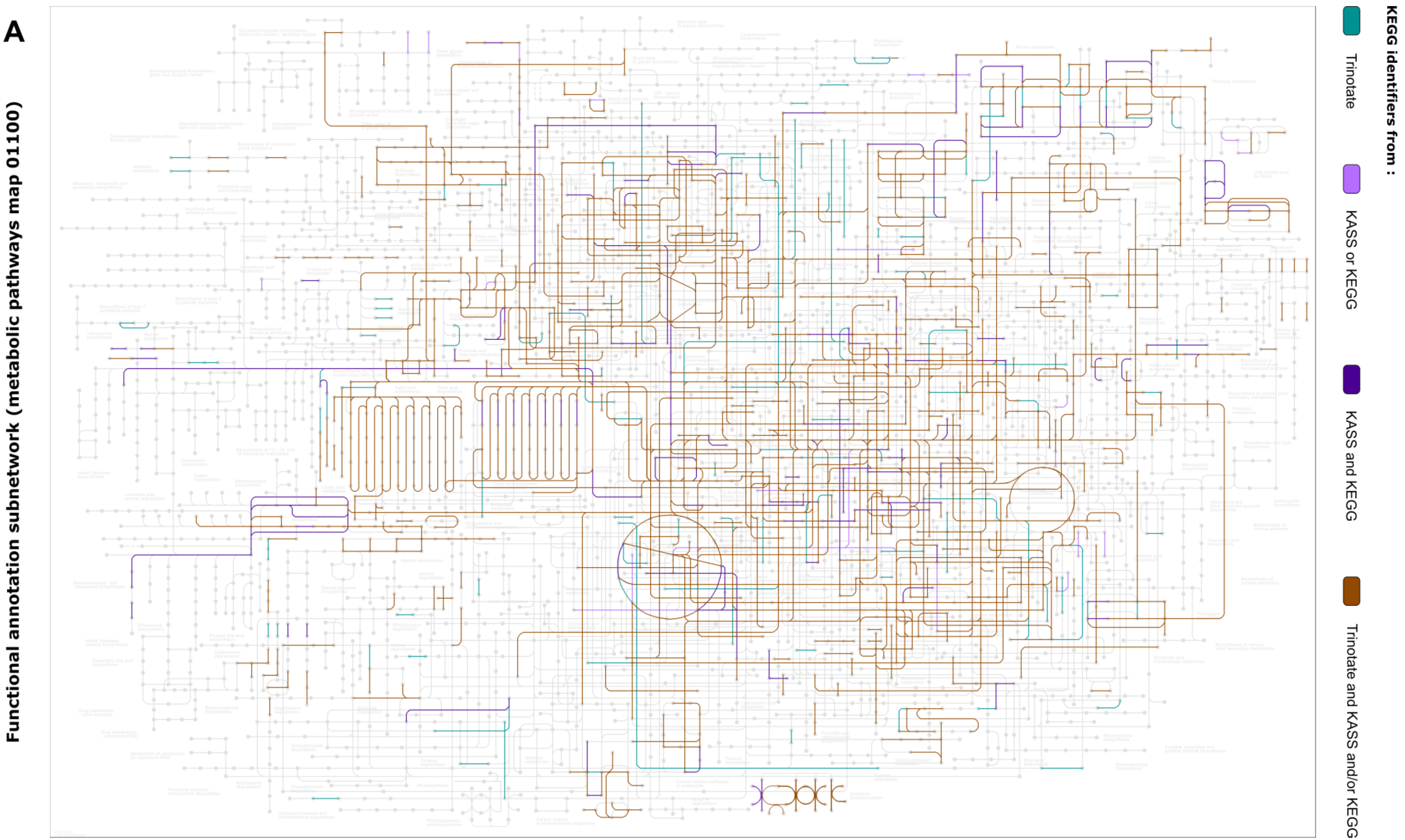

**Figure S2.A: Overview of *Penicillium rubens* reactions performed with KEGG Mapper[1]: complementarity of approaches related to the generation of the functional annotation subnetwork.**

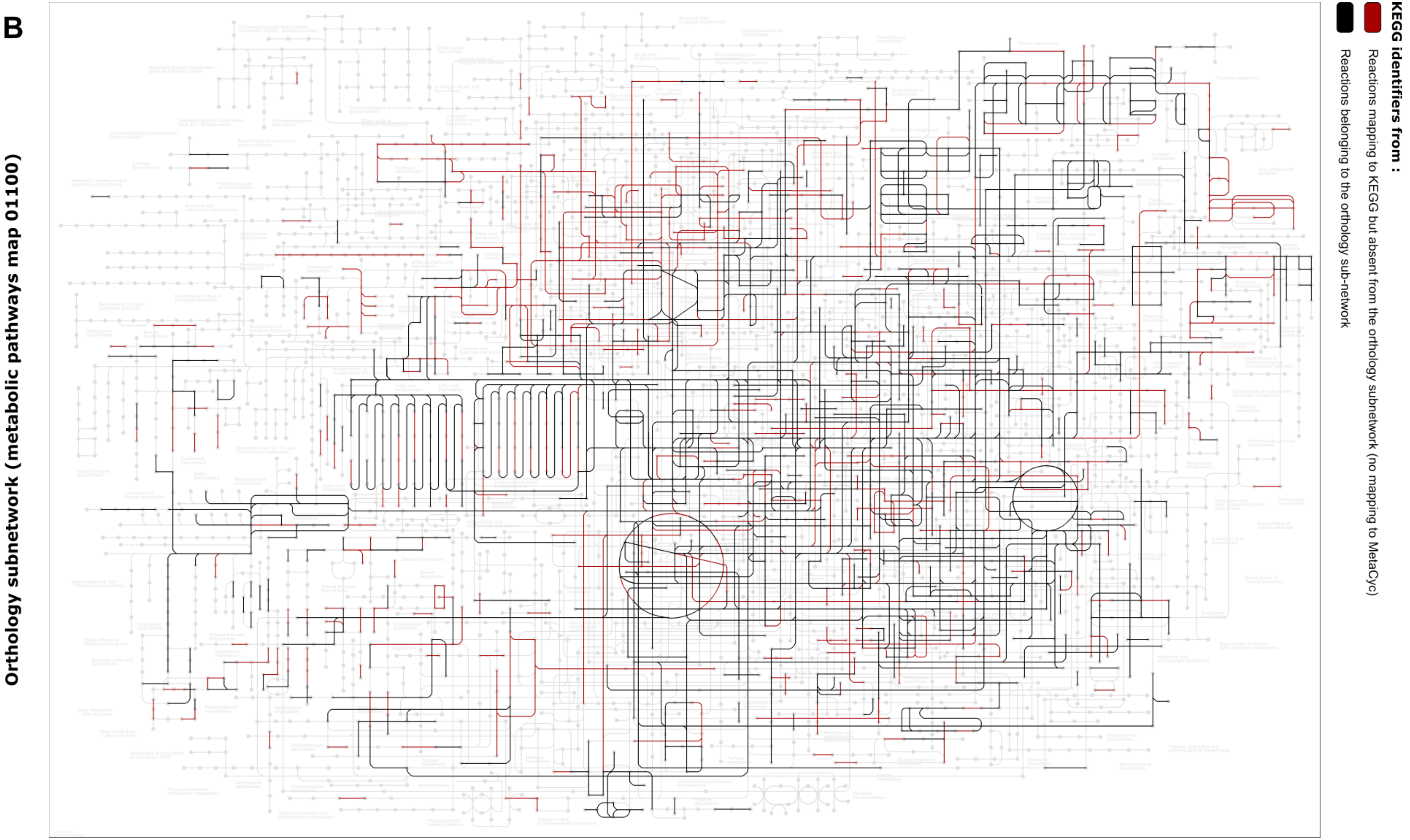

**Figure S2.B: Overview of *Penicillium rubens* reactions performed with KEGG Mapper[1]: losses related to the generation of the orthology subnetwork resulting from mapping operations between databases**

**C**

**Orthology subnetwork (metabolic pathways map 01100)**

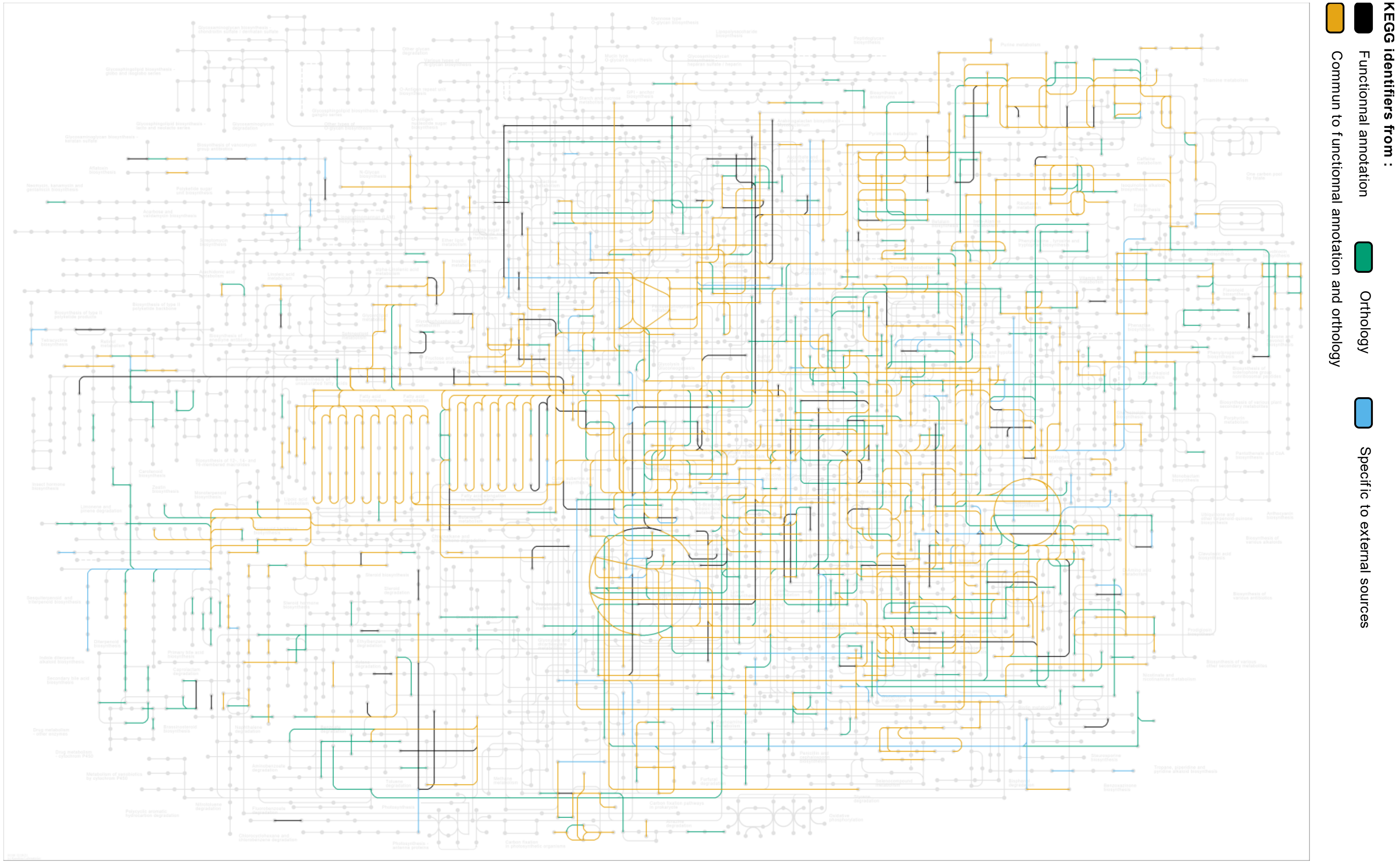

**Figure S2.C: Overview of *Penicillium rubens* reactions performed with KEGG Mapper[1]: topology of the draft after mapping to the KEGG database.**

**Figure S2.A: Overview of *Penicillium rubens* reactions performed with KEGG Mapper:[1] complementarity of approaches related to the generation of the functional annotation subnetwork.** The map is made from the 4,567 KOs found by functional annotation. However, the presence of such an identifier does not guarantee the mapping to a KEGG[2] reaction, let alone a MetaCyc[3] one. The reactions shown were detected *via* the KOs ids found exclusively by Trinotate[4] (■ – 428 KO), KAAS[5] or KEGG (■ – 96 KO), KAAS and KEGG (■ – 378 KO) and Trinotate and KAAS or/and KEGG (■ – 3,665 KO). Although the database query is the same for all three approaches (Trinotate, KAAS and pre-existing *P. rubens* data), there are some slight differences specific to each method.

**Figure S2.B: Overview of *Penicillium rubens* reactions performed with KEGG Mapper:[1] losses related to the generation of the orthology subnetwork resulting from mapping operations between databases.** The map is made from the 2,826 reactions found by orthology search[6]. Representation of reactions belonging to the orthology sub-network (■ – 1,560 reactions) and visualisation of reactions mapping to KEGG but absent from the orthology subnetwork (no mapping to MetaCyc) (■ – 1,266 reactions).

**Figure S2.C: Overview of *Penicillium rubens* reactions performed with KEGG Mapper:[1] topology of the draft after mapping to the KEGG database.** Visualisation of complementarity from functional annotation (■), orthology search (■), shared reactions from both approaches (■), and external sources (■). The various subnetwork reactions are mapped onto the KEGG database using AuReMe[7]. Only 36.57% of the reactions find a match for the annotation sub-network, and this percentage increases to 52.99% for the orthology sub-network. Thus, this visualisation is composed of 228, 627, 933 and 131 reactions (*i.e.* 39.48% of all draft reactions) coming respectively from the annotation subnetwork, the orthology subnetwork, the intersection of these two subnetworks and the external sources.

### *References*

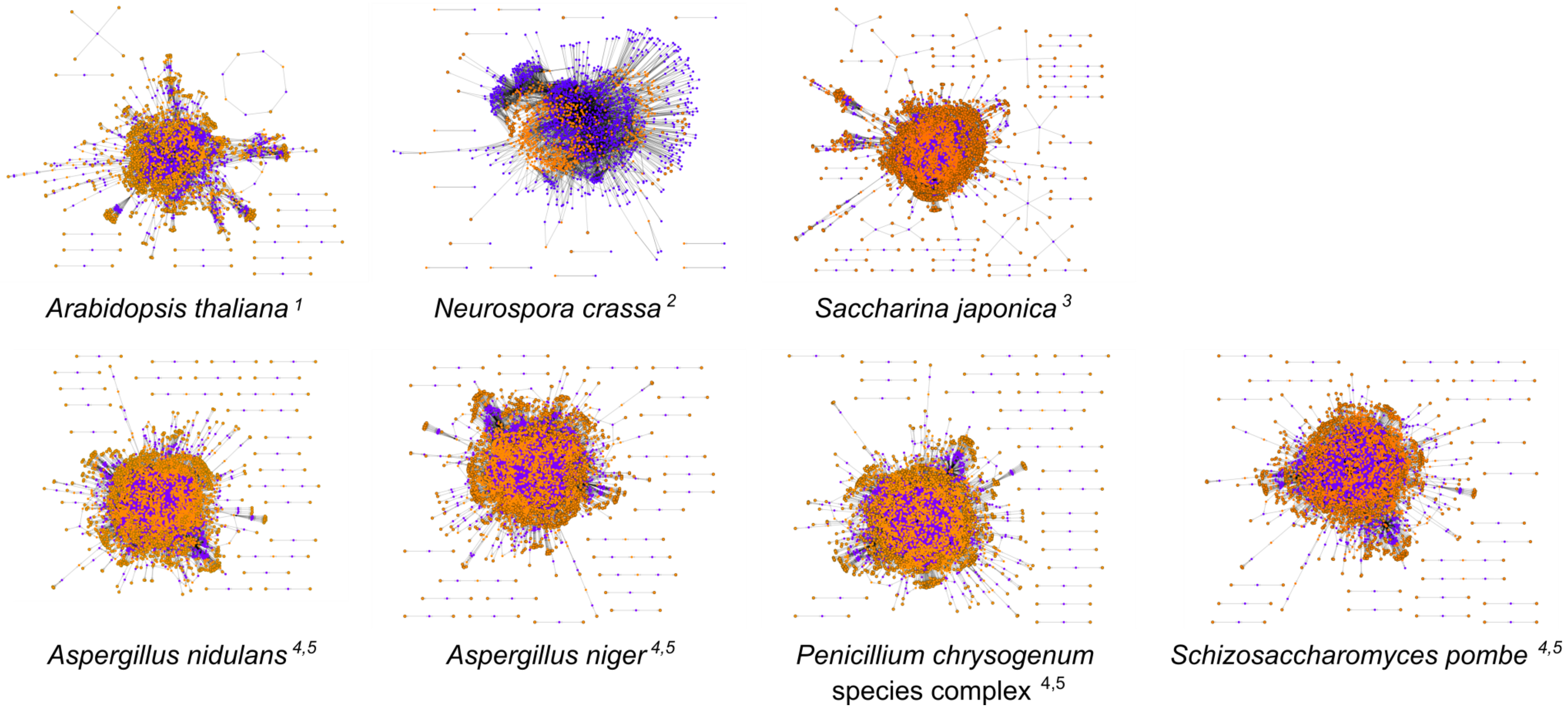

**Figure S3: Bipartite graph representing the seven templates networks' topology for the orthology subnetwork generation.** Reactions and metabolites are shown as purple (■) and orange (■) nodes, respectively. Images obtained with ModelExplorer[6] (version 2.1). GSMNs of *Aspergillus nidulans*, *Aspergillus niger*, *Penicillium rubens* species complex, and *Schizosaccharomyces pombe* are the result of the same automatic reconstruction process. *Saccharina japonica* GSMN has been chosen because its reconstruction process is identical to that presented in this work. In these visualisations, the noticeably higher number of reactions in the *N. crassa* network revealed a defect in the compartmentalisation management.

### A.1 OrthoFinder[1] - Statistics

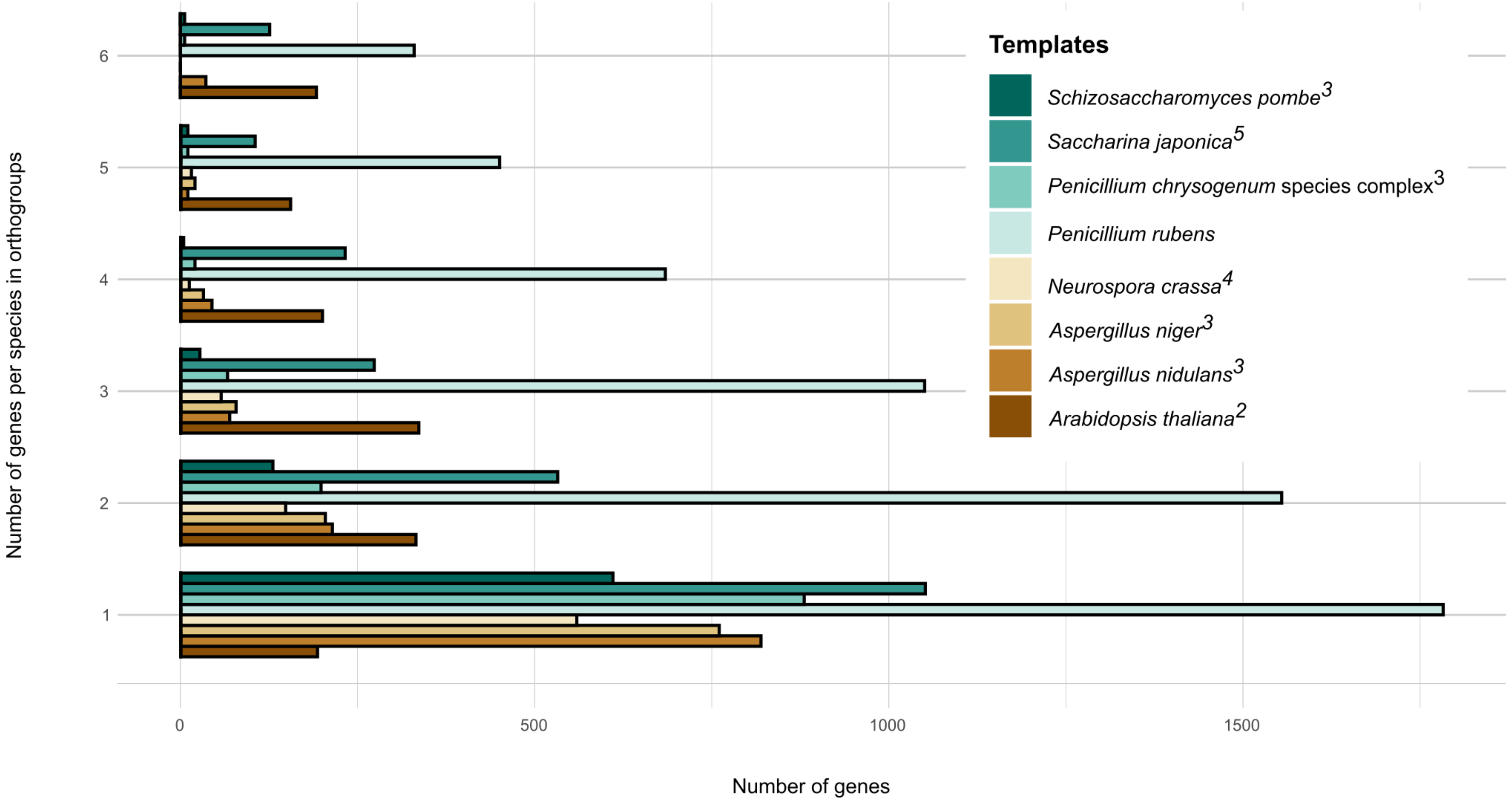

### A.2 OrthoFinder[1] - Statistics

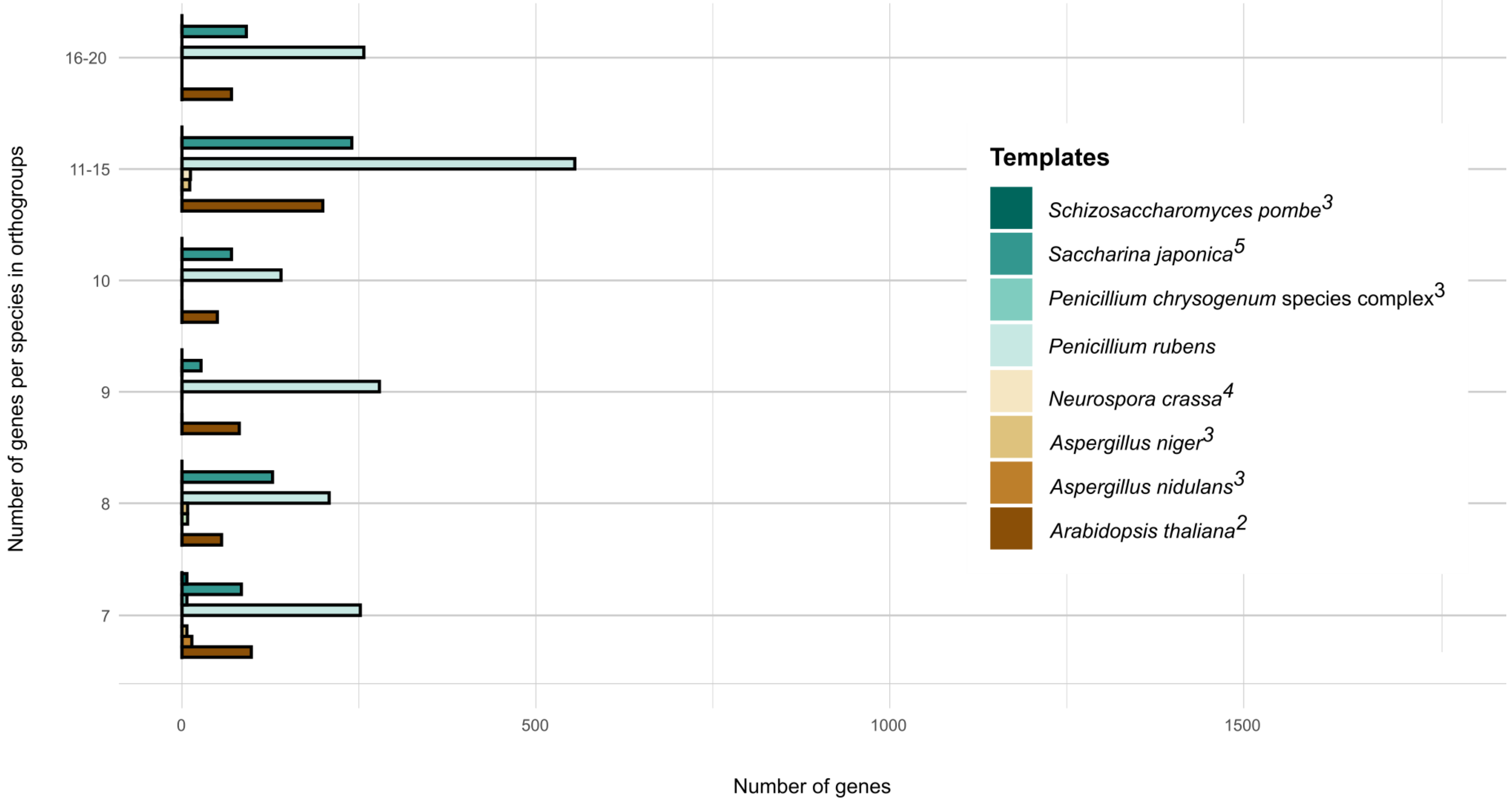

### A.3 OrthoFinder[1] - Statistics

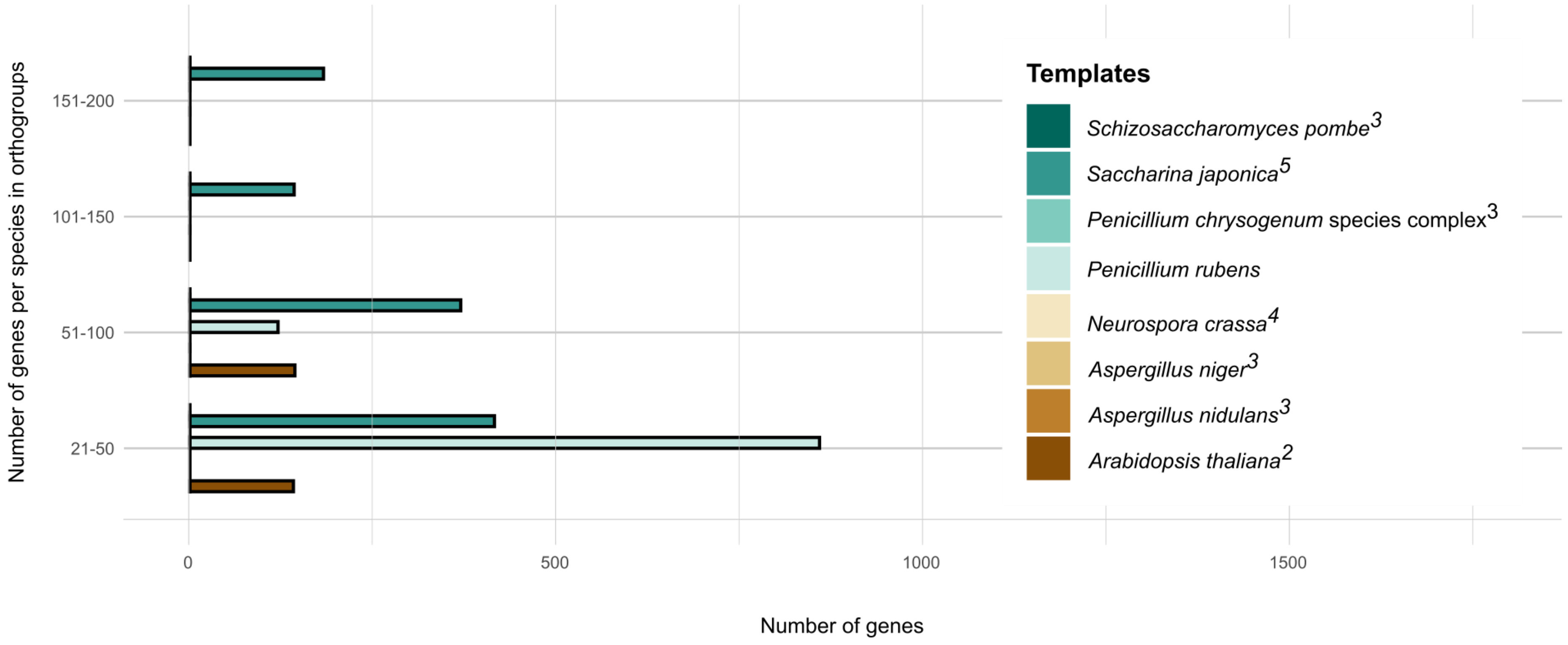

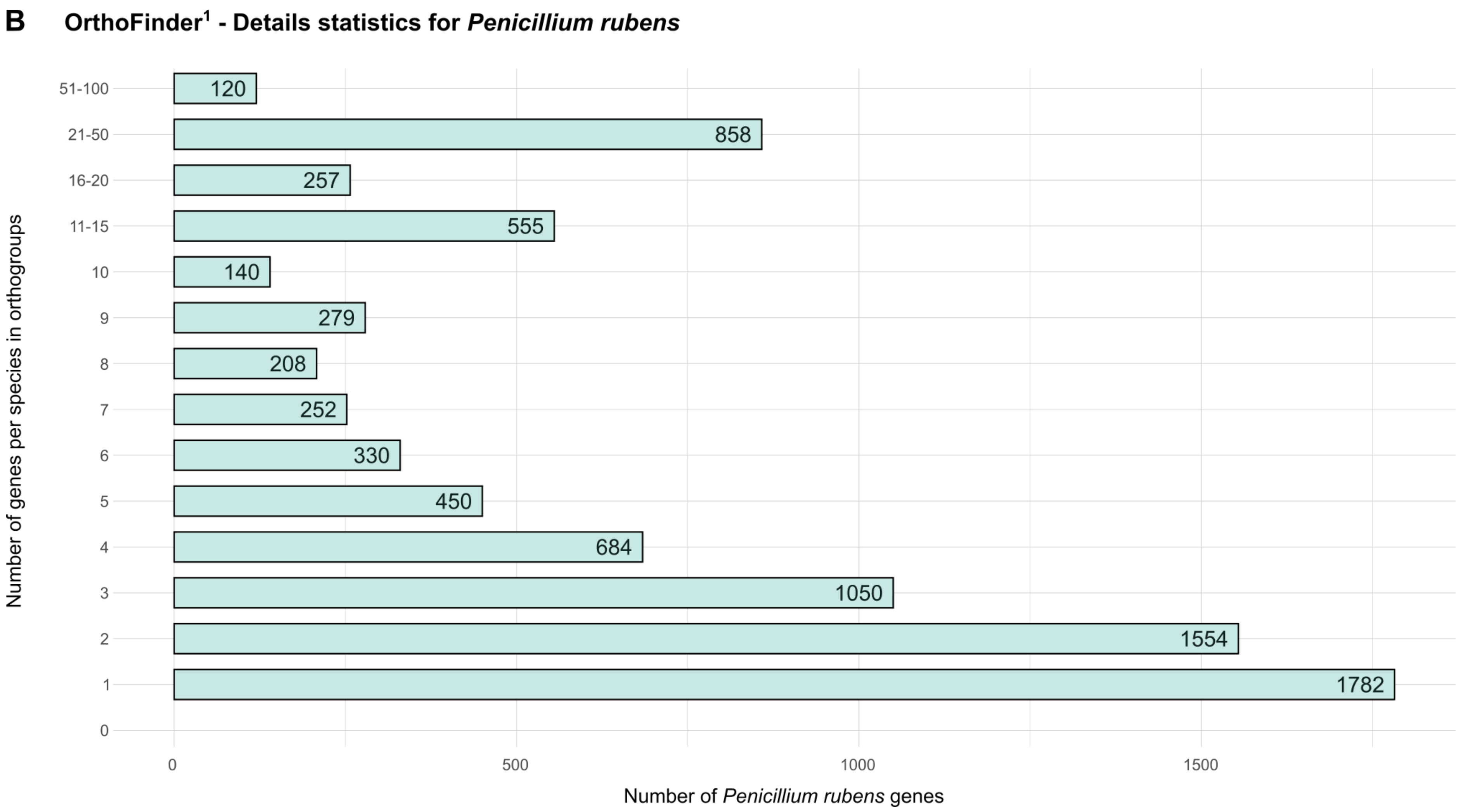

**Figure S4: Sequences number distribution per orthogroup according to species detected with OrthoFinder.**[1] Barplots display this information for all species (A) and then focus on *Penicillium rubens* Wisconsin 54-1255 (B). Whether a pair of genes is an ortholog or a paralogue depends on the event (*i.e.* speciation or duplication) that allows their last common ancestor to be found. Within an orthogroup (a grouping of similar genes derived from a single gene belonging to the closest common ancestor), there is no distinction between these two notions, but their definition is necessary to isolate the orthologous sequences.

### *References*

**A**

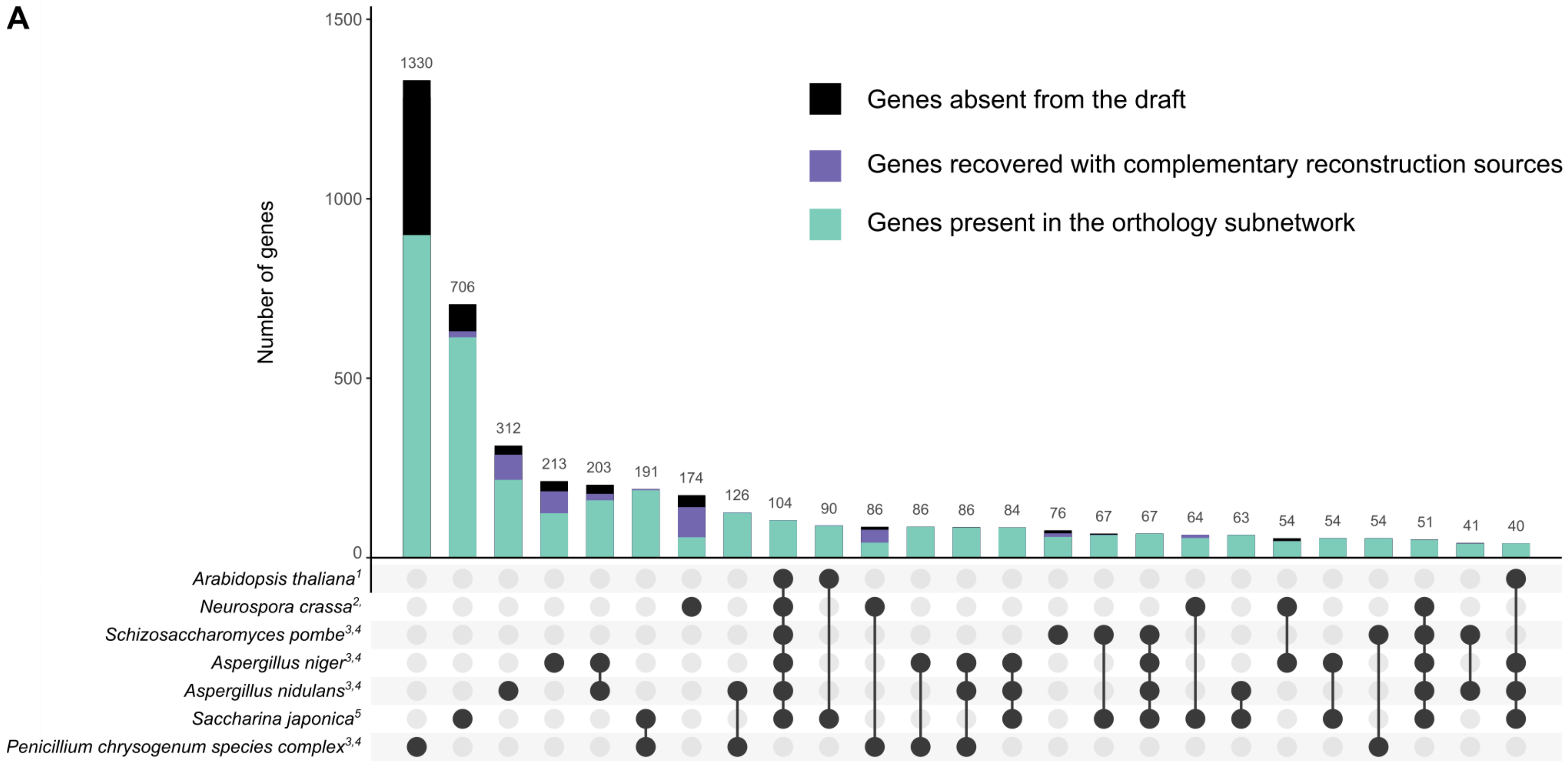

Genes grouping by frequency – Total pool is 5,616 reactions and 4,422 genes are shown on this upset plot
(intersections with a frequency of less than 40 are not displayed)

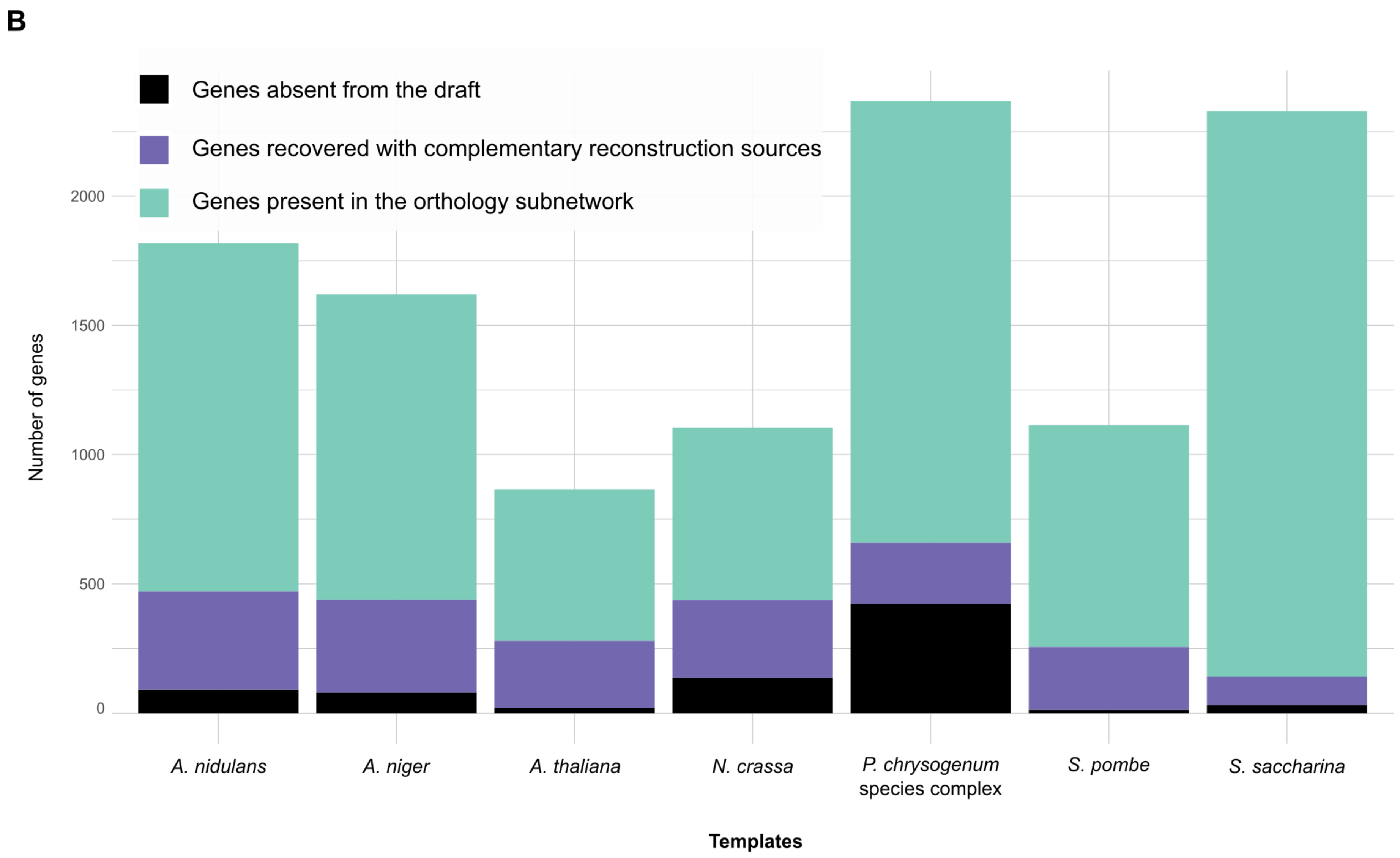
B
Genes absent from the draft
Genes recovered with complementary reconstruction sources
Genes present in the orthology subnetwork
Number of genes
0
500
1000
1500
2000
A. nidulans
A. niger
A. thaliana
N. crassa
P. chrysogenum species complex
S. pombe
S. saccharina
Templates

**C**

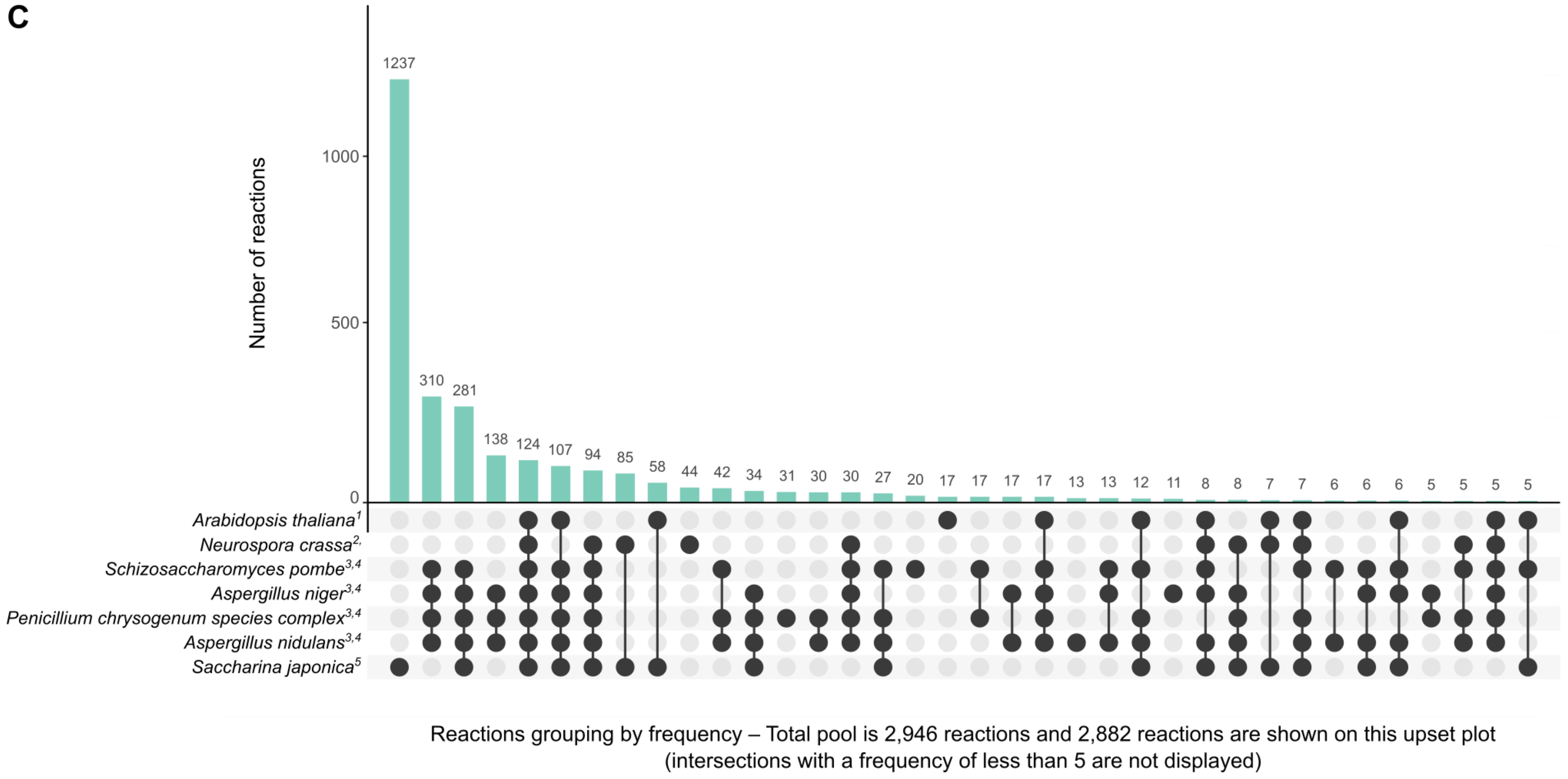

**Figure S5: Visualisation of orthologous genes detected by OrthoFinder and inference of their reactions to the orthology subnetwork. (A)** Upset plot showing the complementarity between templates. Of the 12,556 proteins present in the proteome of the organism studied, 5,616 of them have at least one orthologous sequence in one of the selected templates. Nevertheless, 1,023 of them are not found in the orthology subnetwork due to a lack of interoperability between reaction identifiers. However, 716 of them are then retrieved and added *via* external sources or the functional annotation subnetwork. **(B)** Stacked barplot representing this distribution by the template. **(C)** Upset plot representing the reaction incorporated in the draft according to their orthology source (reactions added after mapping using AuReMe[6]). Figures generated with the R package UpSetR (v1.4.0)[7] and ggplot2 (v3.3.5)[8].

### *References*

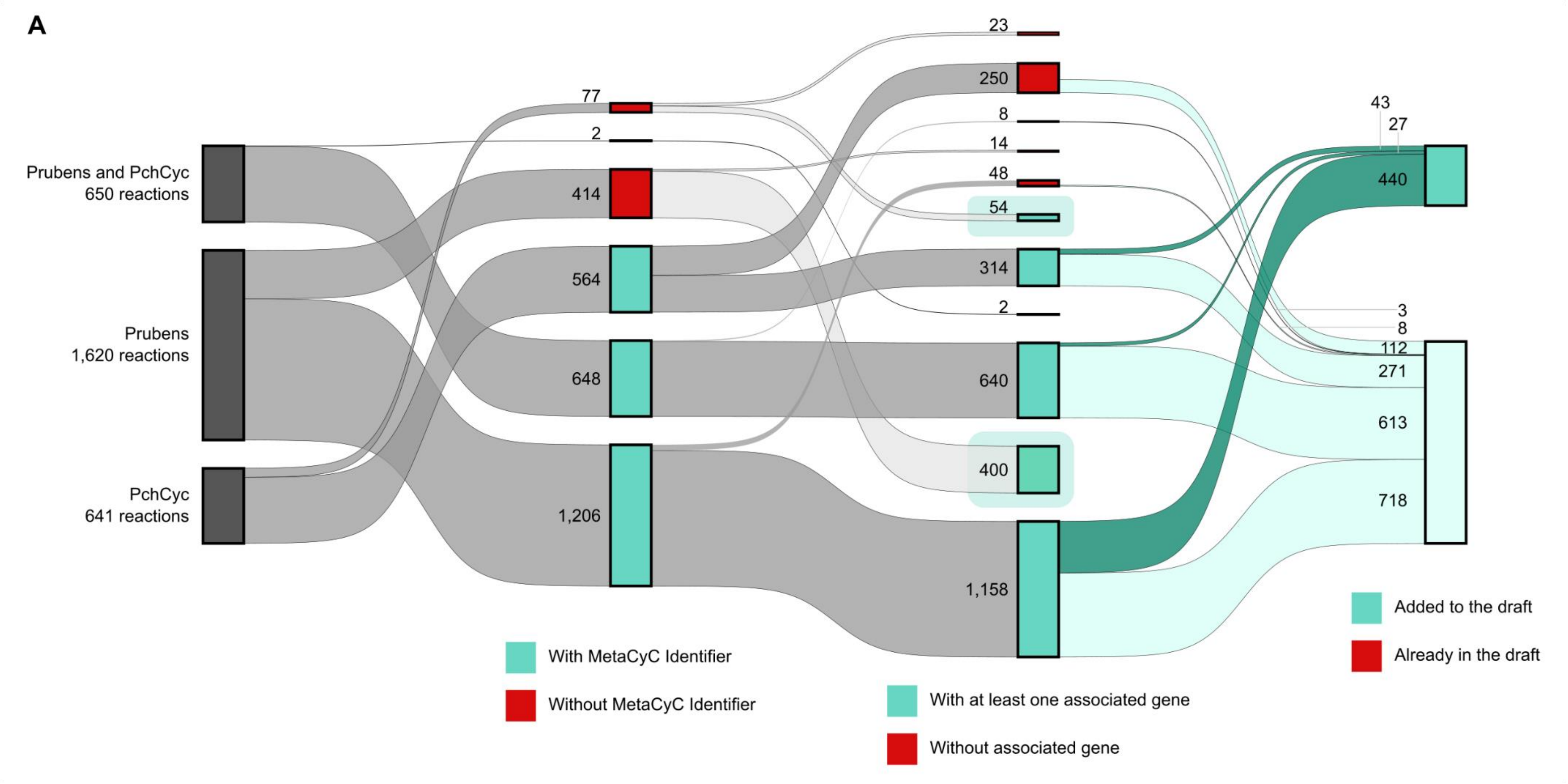
A
Prubens and PchCyc
650 reactions
Prubens
1,620 reactions
PchCyc
641 reactions
77
2
414
564
648
1,206
23
250
8
14
48
54
314
2
640
400
1,158
43
27
440
3
8
112
271
613
718
With MetaCyC Identifier
Without MetaCyC Identifier
With at least one associated gene
Without associated gene
Added to the draft
Already in the draft

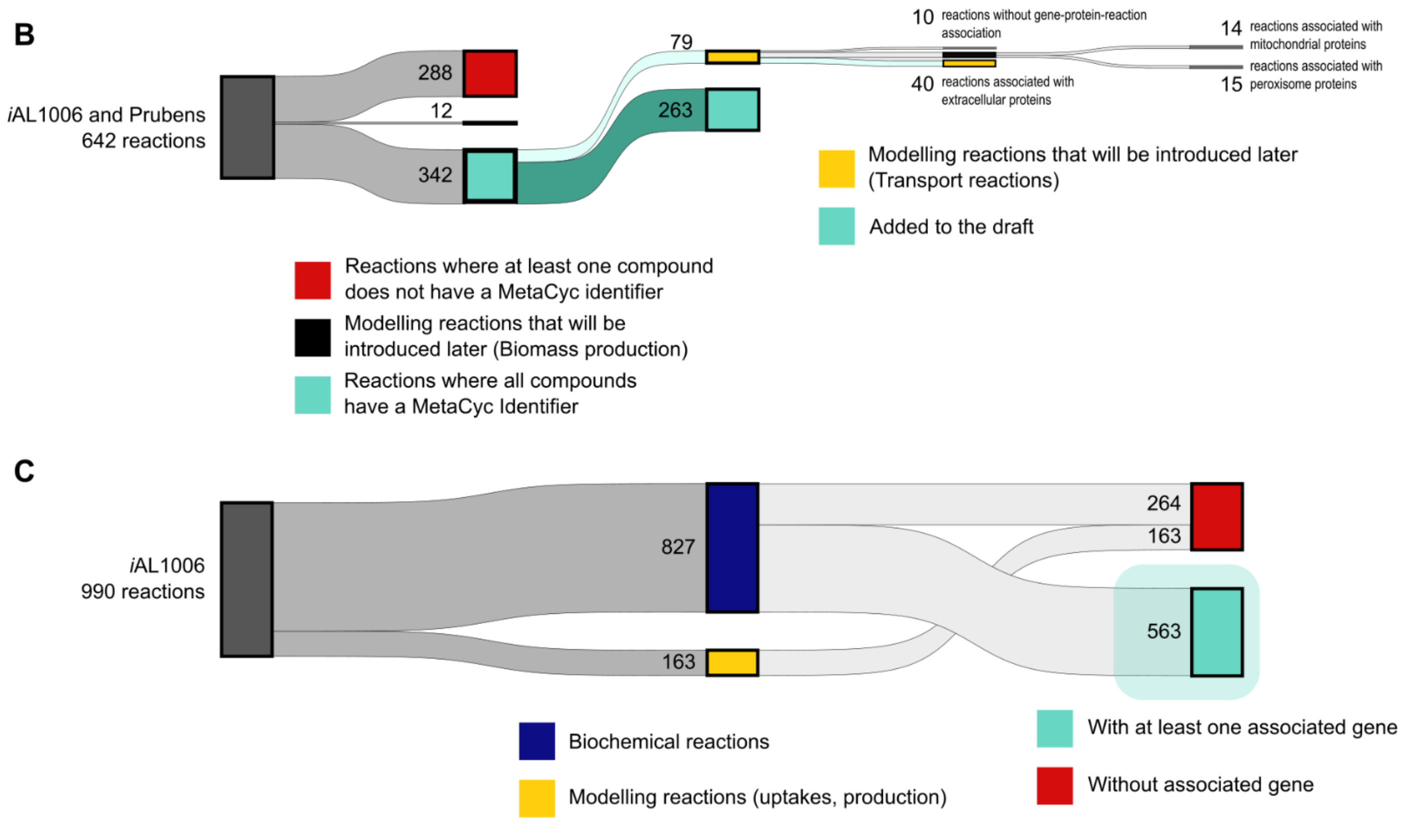

**Figure S6: Sankey plot of the data selection from external sources.** Disregarding compartmentation, the previous *Penicillium rubens* reconstructions are composed of 1,633 reactions for *i*AL1006, 2,533 reactions for Prubens and 1,291 reactions for PchCyc. Reactions were only added to the draft if they had MetaCyc identifiers and were supported by at least one gene (*i.e.* presence of a gene-protein-reaction association - GPR). Of the 4,543 reactions available, only 11% (*i.e.* 510 reactions) were incorporated and 38% (*i.e.* 1,725 reactions) were already present in one of the two automatically-reconstructed subnetworks (*i.e.* source annotation and/or orthology). Finally, it should be noted that 22% of the 4,543 reactions could not be added to this new reconstruction due to the lack of compatible MetaCyc identifiers (*i.e.* 1,017 reactions highlighted by a green rectangle). **(A)** Selection from the most recent data based on MetaCyc-compatible reaction identifiers and the presence of GPRs. **(B)** Selection from reactions present in *i*AL1006 and Prubens based on MetaCyc-compatible compound identifiers and the presence of GPRs. **(C)** Reactions from *i*AL1006 were essentially lost during reconciliation. However, the modelling reactions (*i.e.* uptake and production) were included in the draft in subsequent manual curation steps.

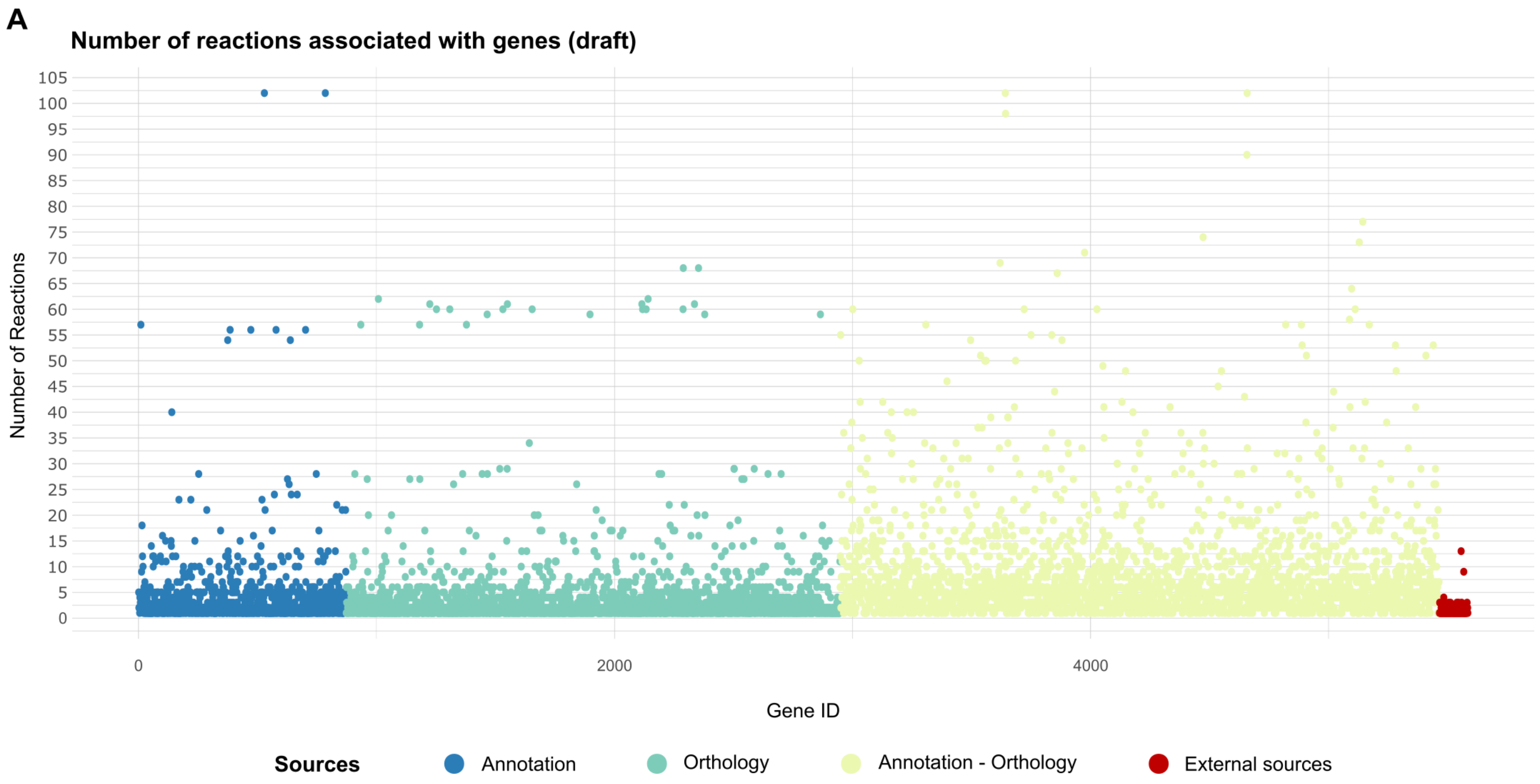
A
Number of reactions associated with genes (draft)
Number of Reactions
105
100
95
90
85
80
75
70
65
60
55
50
45
40
35
30
25
20
15
10
5
0
0
2000
4000
Gene ID
Sources
Annotation
Orthology
Annotation - Orthology
External sources

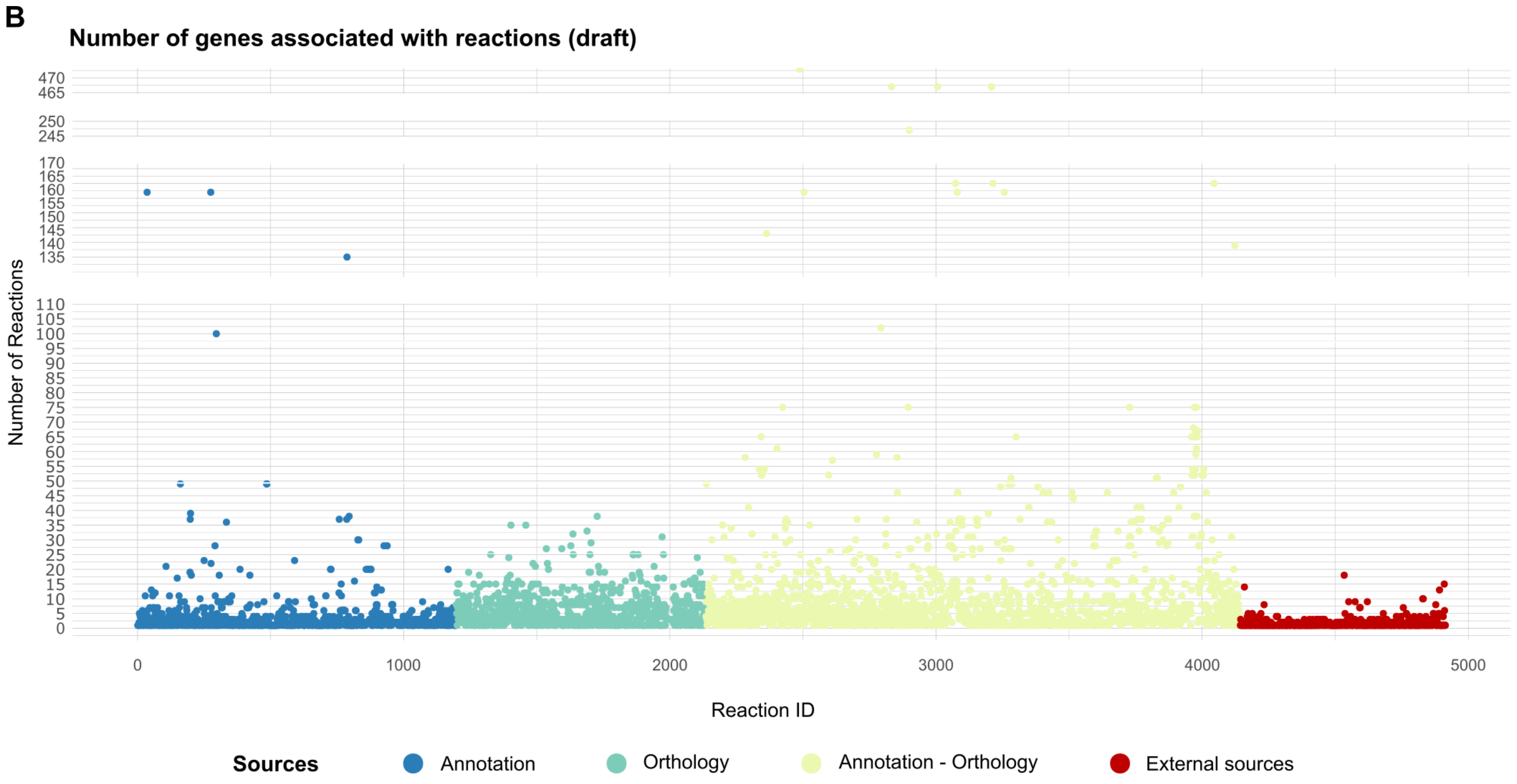

**Figure S7: Scatter plot of reconstruction sources complementarity. (A)** Number of associated reactions per gene and **(B)** number of related genes per reaction. This representation illustrates the complementarity of the various approaches and is an advantage for the further curation of the model. For example, even knowing that GPR associations are not bijective sets, it seems relevant to preferentially check reactions supported exclusively by external sources (■)

**A.1 Classification of genes (subnetwork annotation)**

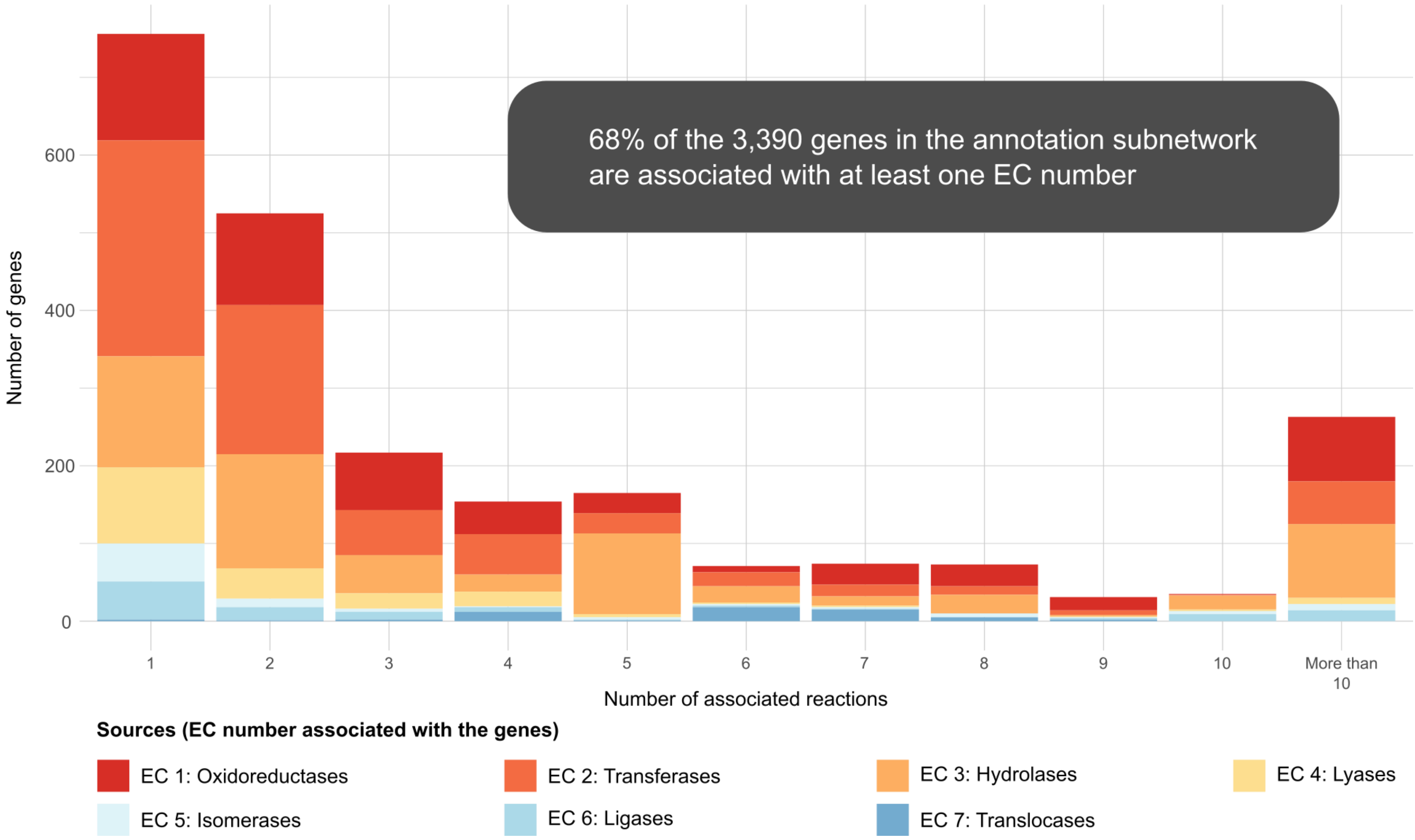

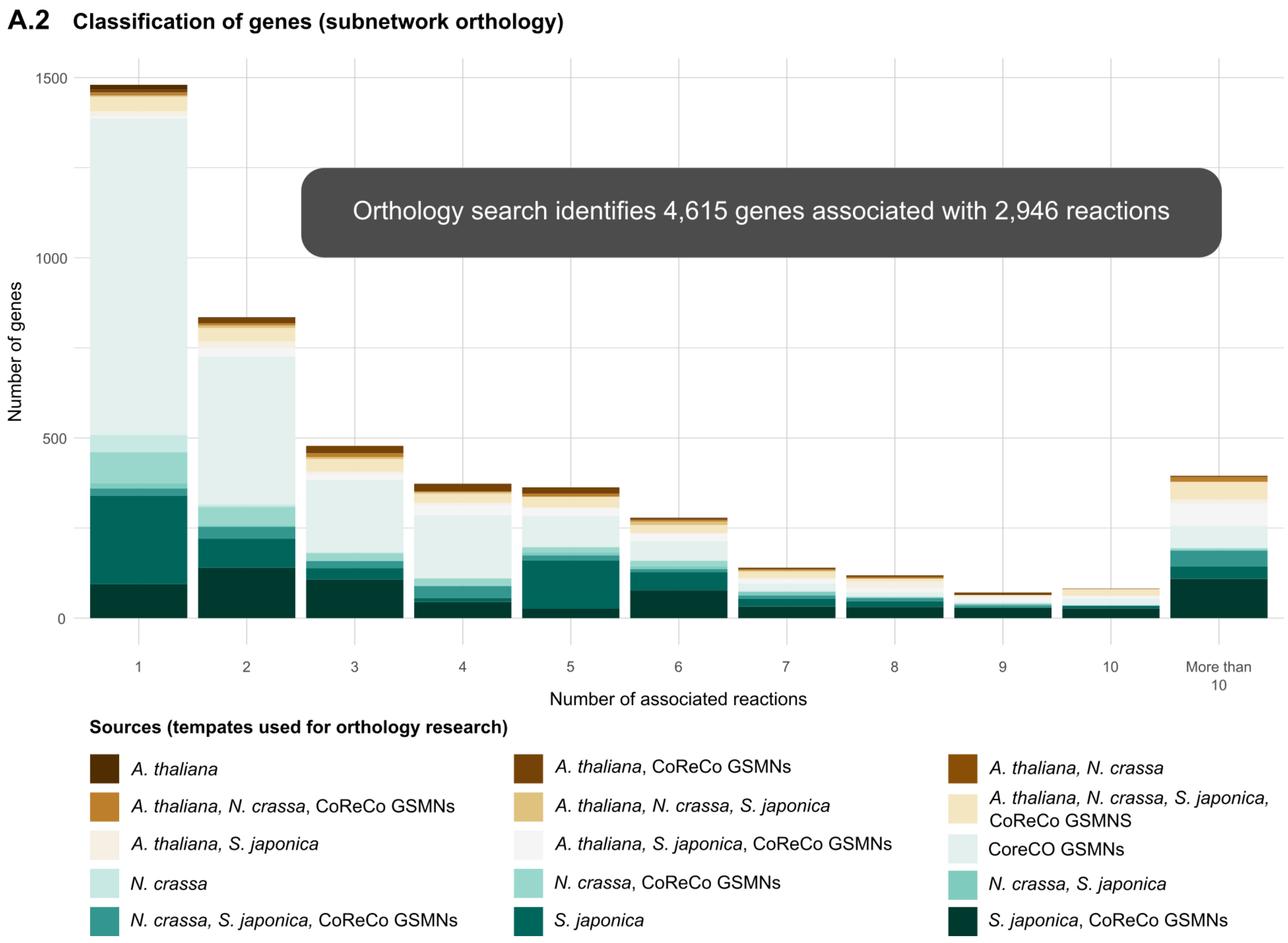
A.2
Classification of genes (subnetwork orthology)
Orthology search identifies 4,615 genes associated with 2,946 reactions
Number of genes
1500
1000
500
0
1
2
3
4
5
6
7
8
9
10
More than 10
Number of associated reactions
Sources (tempates used for orthology research)
A. thaliana
A. thaliana, CoReCo GSMNs
A. thaliana, N. crassa
A. thaliana, N. crassa, CoReCo GSMNs
A. thaliana, N. crassa, S. japonica
A. thaliana, N. crassa, S. japonica, CoReCo GSMNS
A. thaliana, S. japonica
A. thaliana, S. japonica, CoReCo GSMNs
CoreCO GSMNs
N. crassa
N. crassa, CoReCo GSMNs
N. crassa, S. japonica
N. crassa, S. japonica, CoReCo GSMNs
S. japonica
S. japonica, CoReCo GSMNs

**A.3 Classification of genes (draft)**

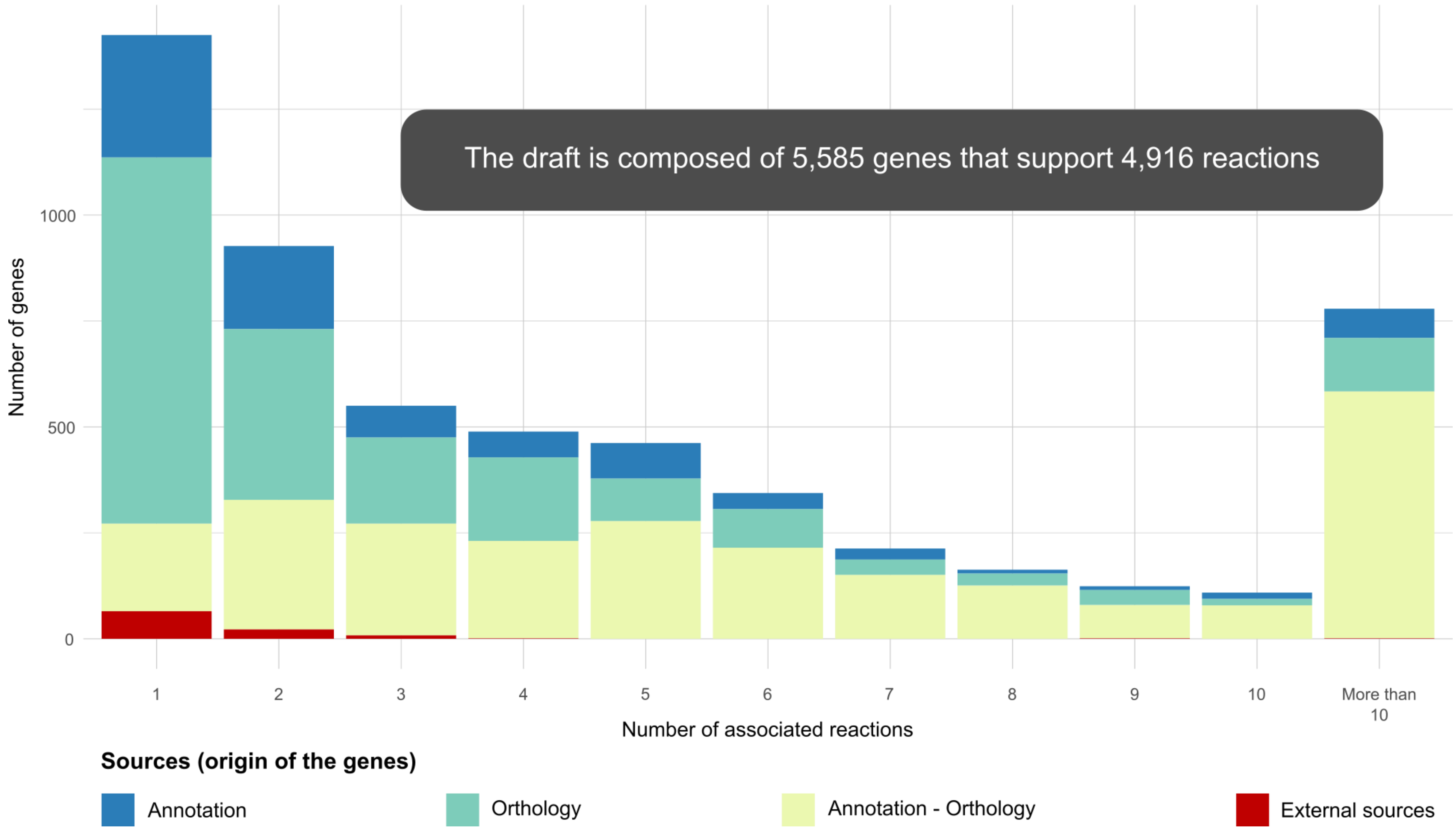

**B.1** **Classification of reactions (subnetwork annotation)**

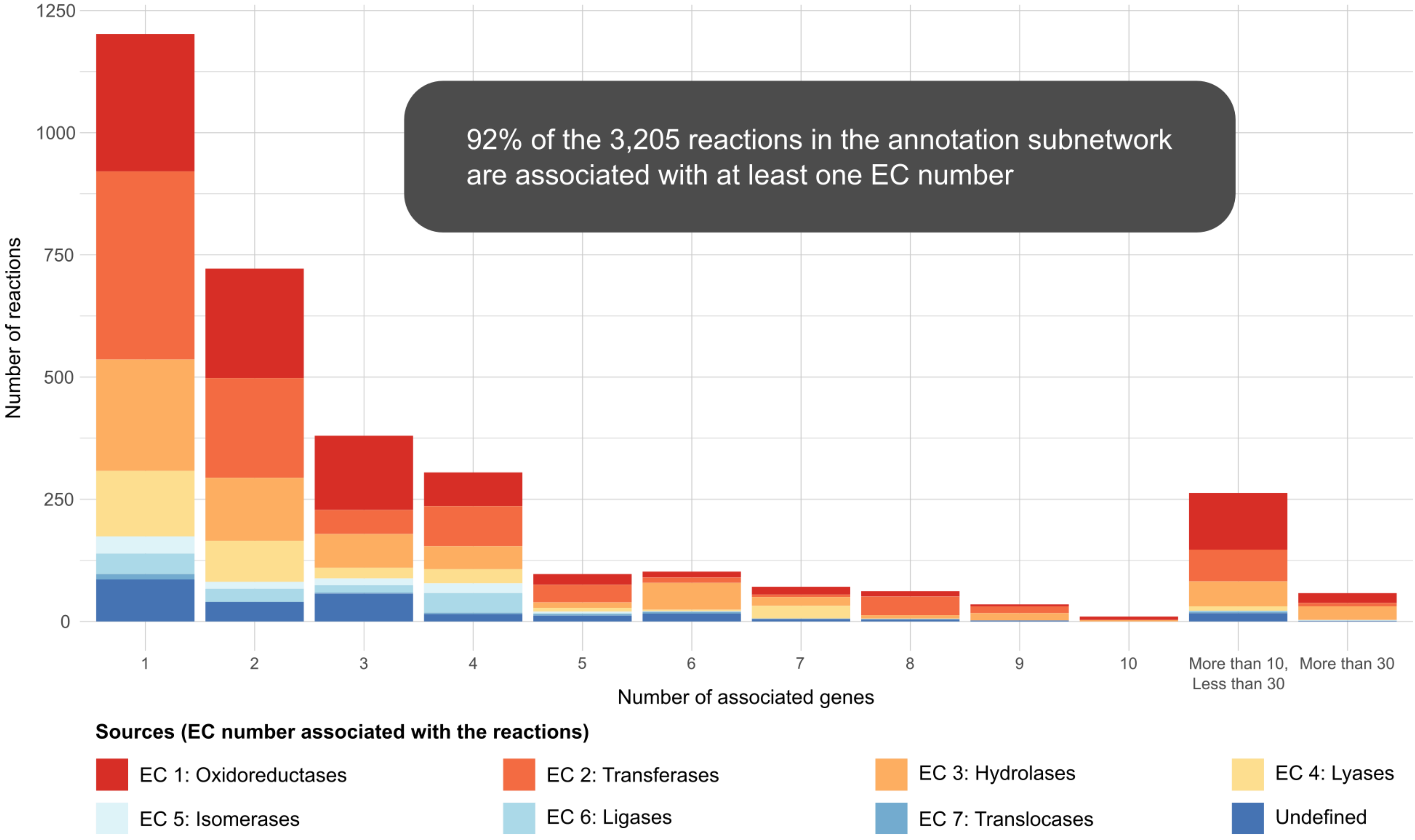

*Supplementary Figure 8 – Results section "The Initial draft" & Discussion section "The strengths and limitations of reconstruction automation"*

**B.2 Classification of reactions (subnetwork orthology)**

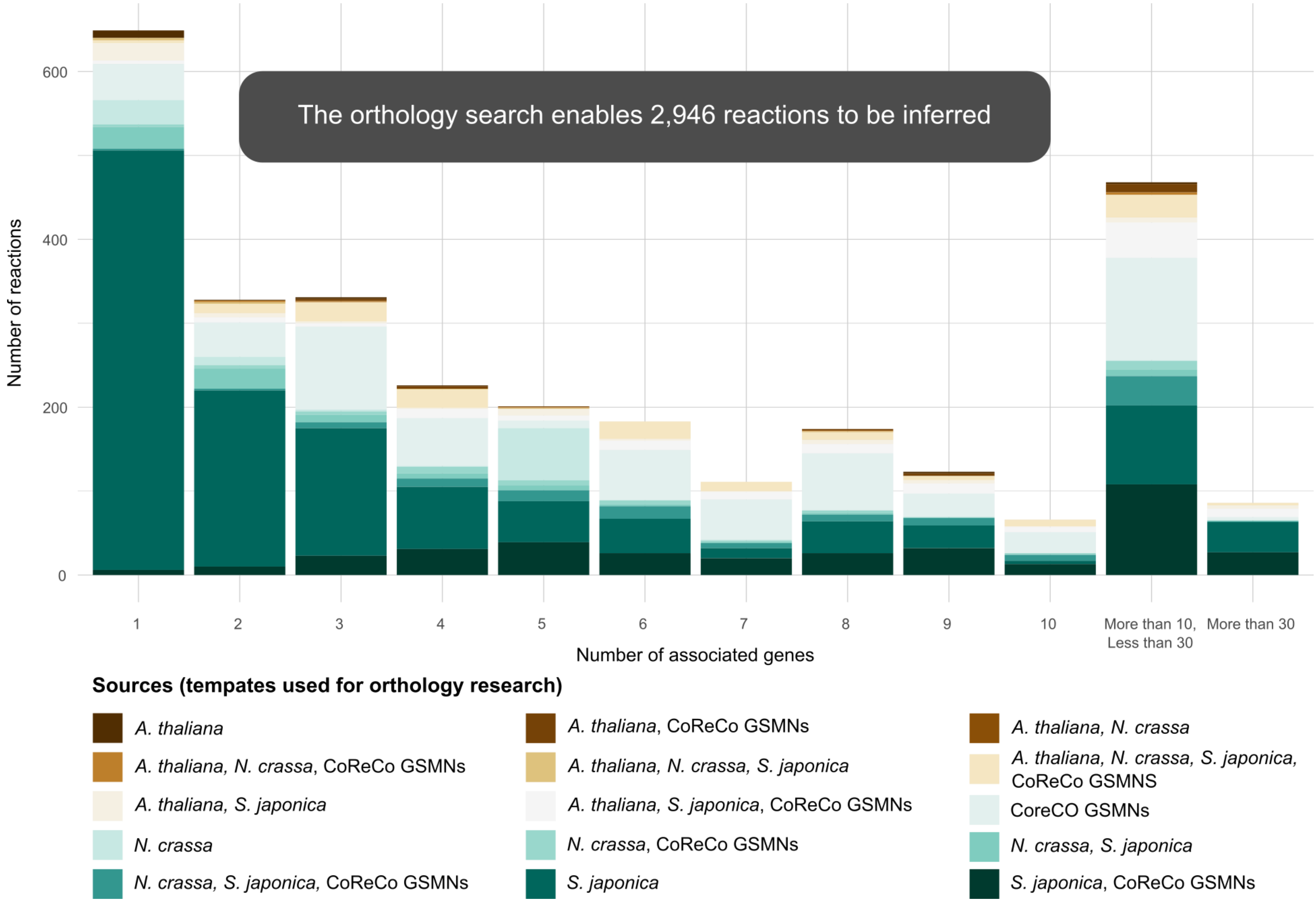

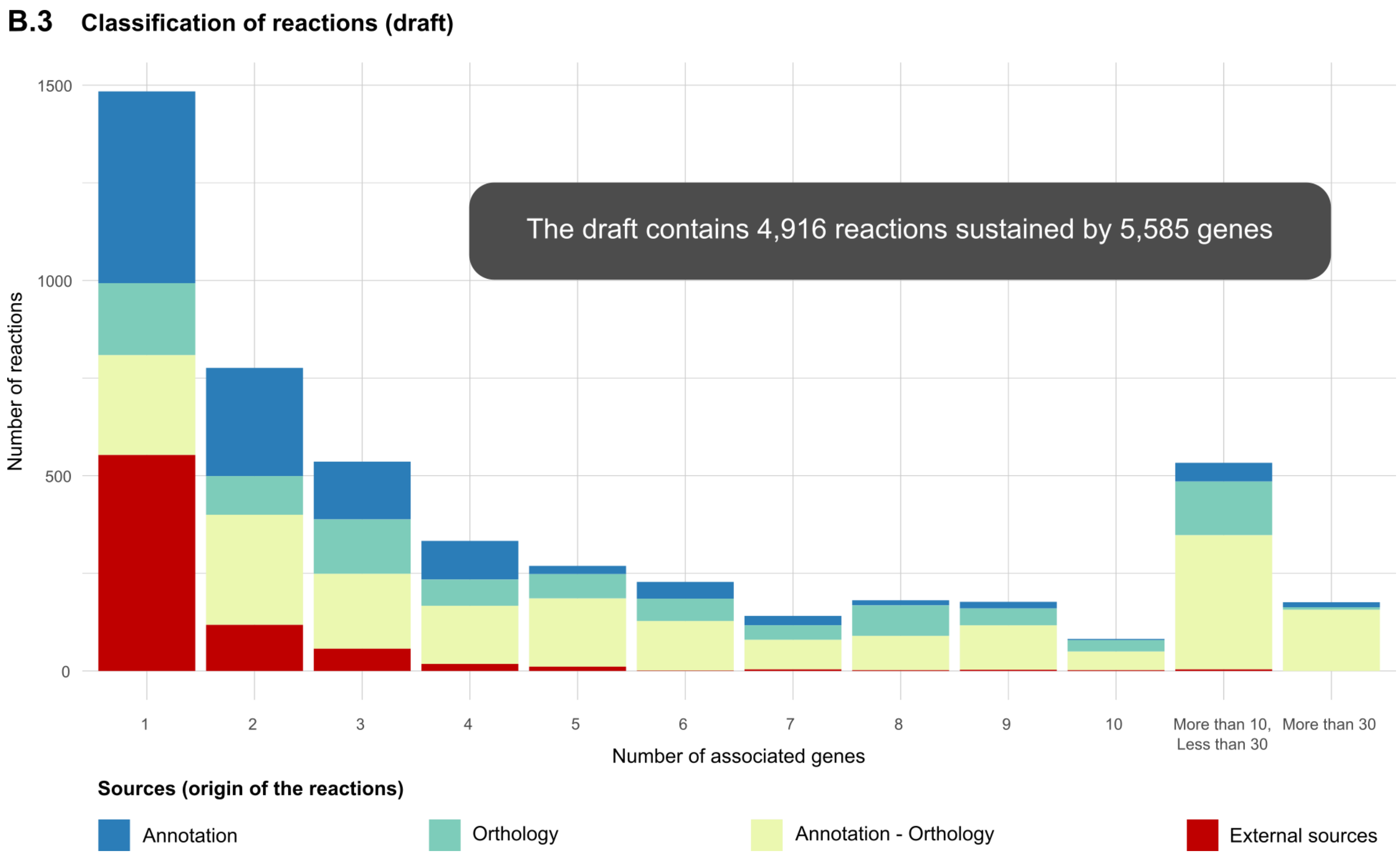

**Figure S8: Classification of genes (A) and reactions (B) according to their source of integration in the draft.**Respectively, the different parts represent the functional annotation sub-network, the orthology sub-network and the fusion of these two sub-networks with the external data. These representations provide details of the contributions of each of the sub-networks and visualise their impact on the presence of genes and reactions within the draft.

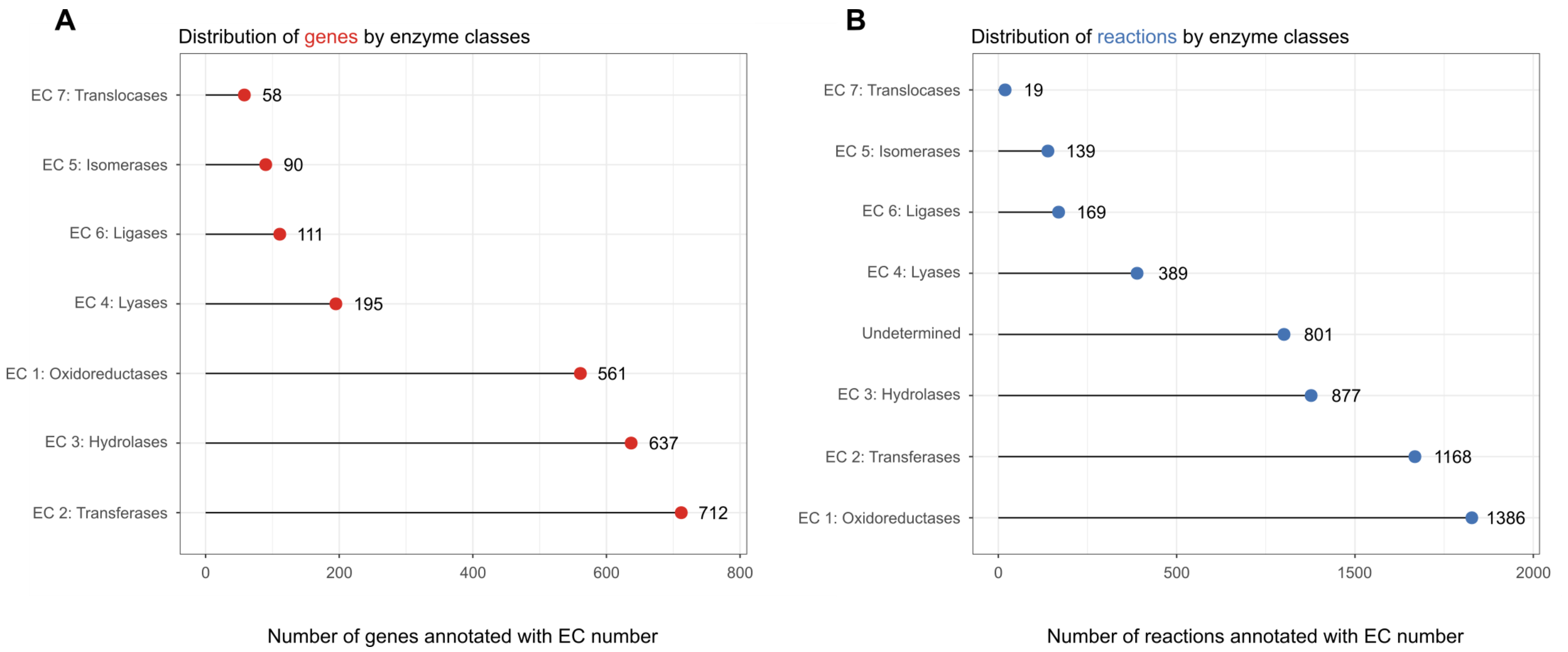

**Figure S9: Classification of genes and reactions according to their associated Enzyme Commission number. (A)** Lollipop plot of genes present in the network and annotated with an EC number mainly during the functional annotation. **(B)** Lollipop plot of reactions according to the EC numbers assigned to them in the GSMN. Respectively 58.42% and 16,29% of genes and reactions lack this type of annotation. Respectively, 219 genes and 1,032 reactions are associated with more than two EC numbers and 42 and 28 of these correspond to at least two different classes. Transferases, hydrolases and oxidoreductases are the most frequent annotations for genes, while oxidoreductases, transfer enzymes, and hydrolases are the most common annotations for reactions. The ratios between the number of genes and reactions are, therefore, variable depending on the enzyme class: oxidoreductases (661:1386); transferases (89:146); hydrolases (637:877); lyases (195:389); isomerases (90:139); ligases (111:169); translocases (58:19).

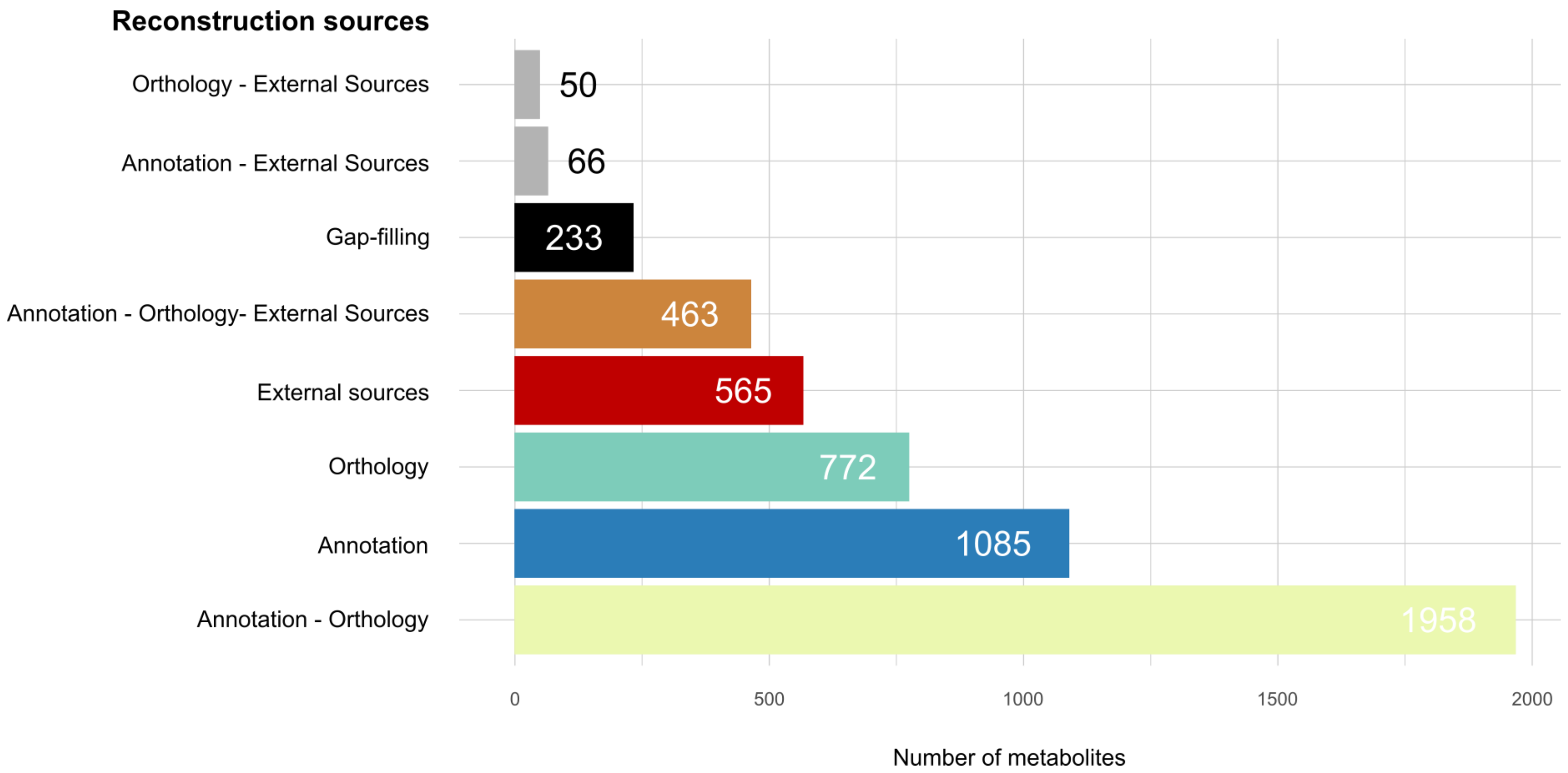

**Figure S10: Origin of metabolites in *i*Prub22 according to reaction reconstruction sources.** Manual curation (gap-filling) will add 232 metabolites to the model.

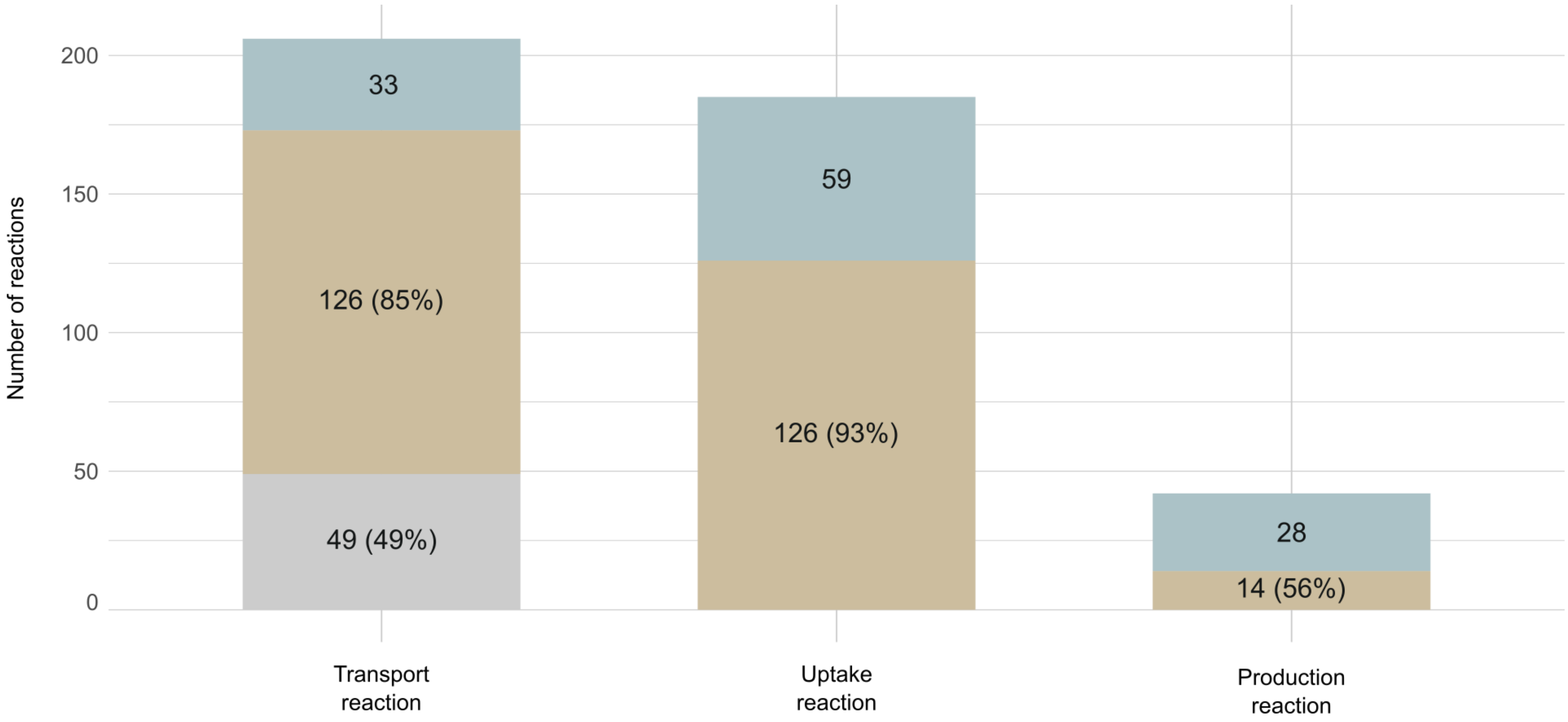

**Figure S11: Sources of transport and exchange reactions added to the reconstruction.** Reactions from *i*AL1006 (■), from PchCyc (■) and for media simulation and/or consistency and/or monitoring specialised metabolites production(■). The percentages in brackets correspond to the reaction number selected from the external sources (*i*AL1006 and PchCyc). The reaction rejection is explained either (1) by the absence of the targeted metabolite within the draft topology, (2) by the absence of a metabolite identification on the MetaCyc database, or, when it exists, (3) by a produced protein annotation location different from the cell membrane. For further details please refer to *Transport_Exchange.xlsx*.

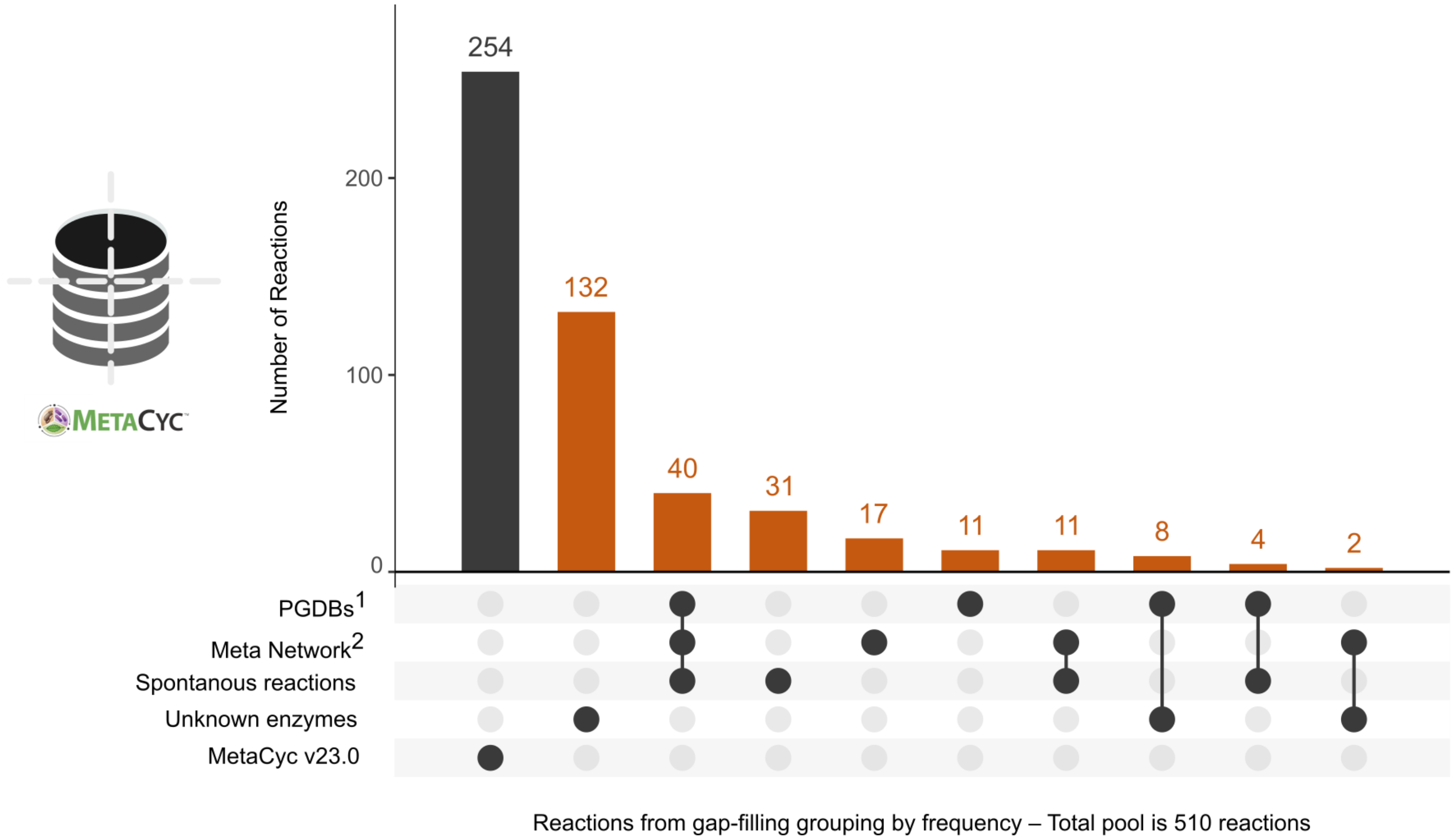

**Figure S12: Distribution of the 510 reactions added to the reconstruction during the gap-filling steps.** The 256 orange reactions (■) detected with MENECO[3] come from the selected subsets of MetaCyc. For half of the reactions added, this "reasoned" gap-filling increases the presence confidence of in these reactions in the reconstruction. For further details please refer to *All_curation.xlsx*. Figure generated with the R package UpSetR (v1.4.0)[4].

### *References*

(1) Caspi, R.; Billington, R.; Keseler, I. M.; Kothari, A.; Krummenacker, M.; Midford, P. E.; Ong, W. K.; Paley, S.; Subhraveti, P.; Karp, P. D. The MetaCyc Database of Metabolic Pathways and Enzymes - a 2019 Update. Nucleic Acids Res. **2020**, 48 (D1), D445–D453. https://doi.org/10.1093/nar/gkz862.

(2) Belcour, A.; Got, J.; Aite, M.; Delage, L.; Collen, J.; Frioux, C.; Leblanc, C.; Dittami, S. M.; Blanquart, S.; Markov, G. V.; Siegel, A. AuCoMe: Inferring and Comparing Metabolisms across Heterogeneous Sets of Annotated Genomes. bioRxiv June 17, **2022**, p 2022.06.14.496215. https://doi.org/10.1101/2022.06.14.496215.

(3) Prigent, S.; Frioux, C.; Dittami, S. M.; Thiele, S.; Larhlimi, A.; Collet, G.; Gutknecht, F.; Got, J.; Eveillard, D.; Bourdon, J.; Plewniak, F.; Tonon, T.; Siegel, A. Meneco, a Topology-Based Gap-Filling Tool Applicable to Degraded Genome-Wide Metabolic Networks. PLOS Comput. Biol. **2017**, 13 (1), e1005276. https://doi.org/10.1371/journal.pcbi.1005276.

(4) Conway, J. R.; Lex, A.; Gehlenborg, N. UpSetR: An R Package for the Visualization of Intersecting Sets and Their Properties. Bioinforma. Oxf. Engl. **2017**, 33 (18), 2938–2940. https://doi.org/10.1093/bioinformatics/btx364.

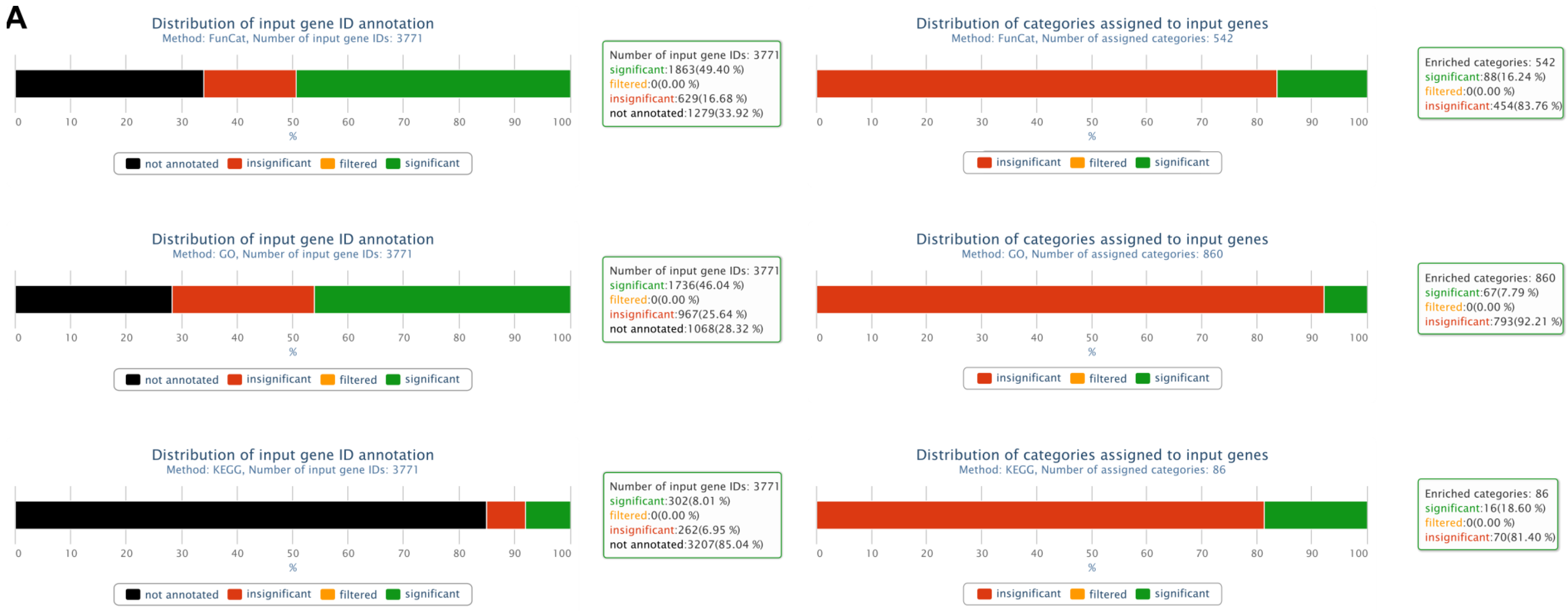
A
Distribution of input gene ID annotation
Method: FunCat, Number of input gene IDs: 3771
Number of input gene IDs: 3771
significant:1863(49.40 %)
filtered:0(0.00 %)
insignificant:629(16.68 %)
not annotated:1279(33.92 %)
not annotated
insignificant
filtered
significant
Distribution of categories assigned to input genes
Method: FunCat, Number of assigned categories: 542
Enriched categories: 542
significant:88(16.24 %)
filtered:0(0.00 %)
insignificant:454(83.76 %)
Distribution of input gene ID annotation
Method: GO, Number of input gene IDs: 3771
Number of input gene IDs: 3771
significant:1736(46.04 %)
filtered:0(0.00 %)
insignificant:967(25.64 %)
not annotated:1068(28.32 %)
Distribution of categories assigned to input genes
Method: GO, Number of assigned categories: 860
Enriched categories: 860
significant:67(7.79 %)
filtered:0(0.00 %)
insignificant:793(92.21 %)
Distribution of input gene ID annotation
Method: KEGG, Number of input gene IDs: 3771
Number of input gene IDs: 3771
significant:302(8.01 %)
filtered:0(0.00 %)
insignificant:262(6.95 %)
not annotated:3207(85.04 %)
Distribution of categories assigned to input genes
Method: KEGG, Number of assigned categories: 86
Enriched categories: 86
significant:16(18.60 %)
filtered:0(0.00 %)
insignificant:70(81.40 %)
%

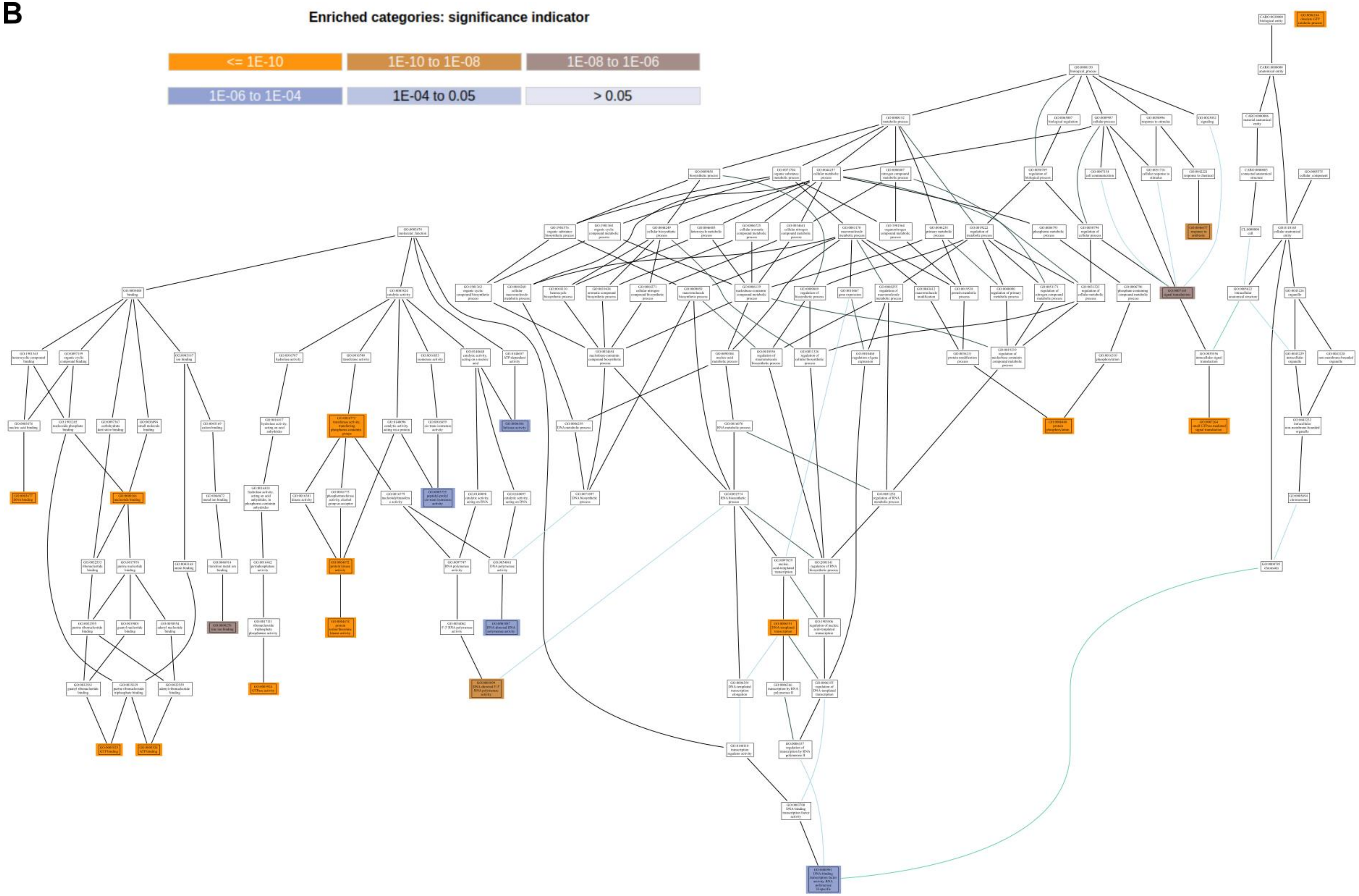
B
Enriched categories: significance indicator
<= 1E-10
1E-10 to 1E-08
1E-08 to 1E-06
1E-06 to 1E-04
1E-04 to 0.05
> 0.05

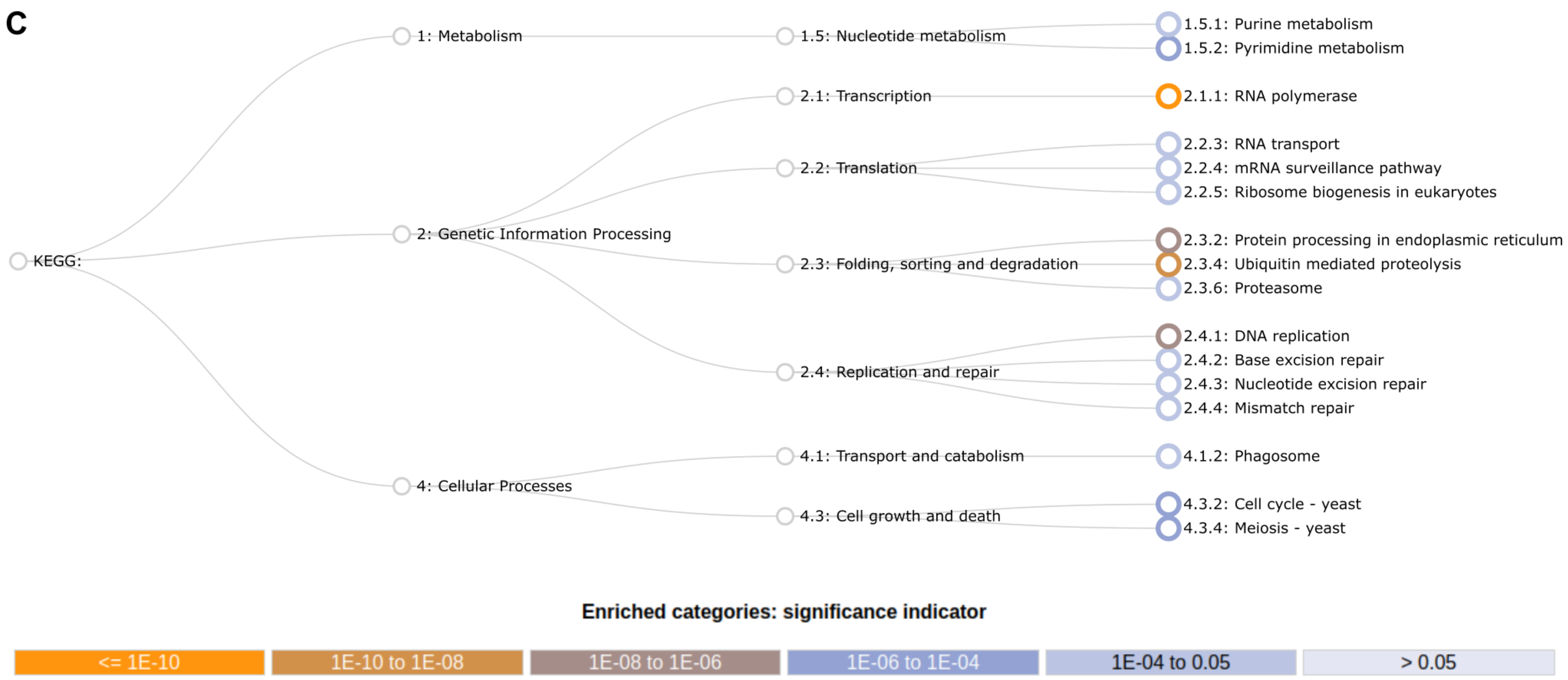
C
KEGG:
1: Metabolism
2: Genetic Information Processing
4: Cellular Processes
1.5: Nucleotide metabolism
2.1: Transcription
2.2: Translation
2.3: Folding, sorting and degradation
2.4: Replication and repair
4.1: Transport and catabolism
4.3: Cell growth and death
1.5.1: Purine metabolism
1.5.2: Pyrimidine metabolism
2.1.1: RNA polymerase
2.2.3: RNA transport
2.2.4: mRNA surveillance pathway
2.2.5: Ribosome biogenesis in eukaryotes
2.3.2: Protein processing in endoplasmic reticulum
2.3.4: Ubiquitin mediated proteolysis
2.3.6: Proteasome
2.4.1: DNA replication
2.4.2: Base excision repair
2.4.3: Nucleotide excision repair
2.4.4: Mismatch repair
4.1.2: Phagosome
4.3.2: Cell cycle - yeast
4.3.4: Meiosis - yeast
Enriched categories: significance indicator
<= 1E-10
1E-10 to 1E-08
1E-08 to 1E-06
1E-06 to 1E-04
1E-04 to 0.05
> 0.05

**D**

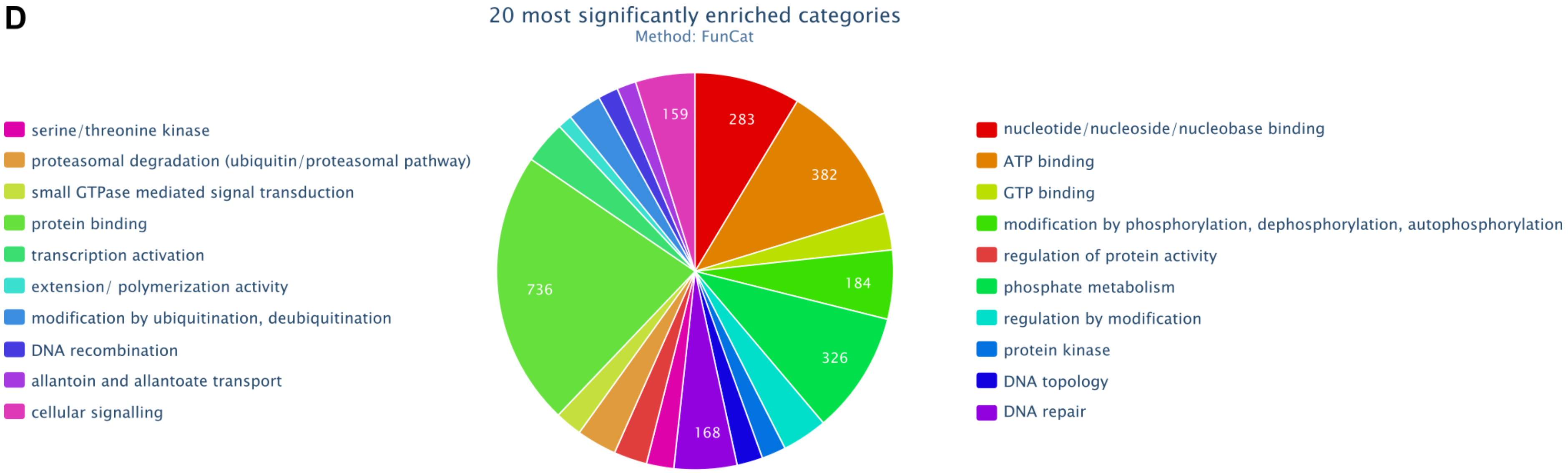

**Figure S13: Annotations enrichment of the 3,771 genes added to the *Penicillium rubens* GSMN reconstruction (Results from FungiFun). (A)** Distribution visualisation of the genes annotations and their categories according to the following ontologies **(B)** GOT, **(C)** KEGG, **(D)** FunCat. The classification is represented by a pie chart or by a hierarchy graph. They show the most significant categories for each queried ontology.

### *Reference*

(1) Priebe, S.; Kreisel, C.; Horn F.; Guthke, R.; Linde, J. FUNGIGUN2: A Comprehensive Online Resource for Systematic Analysis of Gene Lists from Fungal Species. Bioinformatics 2015, 31 (3), 445-446. https://doi.org/10.1093/BIOINFORMATICS/BTU627.
